# Allele-specific gene editing approach for vision loss restoration in *RHO*-associated Retinitis Pigmentosa

**DOI:** 10.1101/2022.11.16.516784

**Authors:** Xiaozhen Liu, Jing Qiao, Ruixuan Jia, Fan Zhang, Xiang Meng, Yang Li, Liping Yang

**Author notes:** Liping Yang, MD, PhD, Room 721, 49 North Garden Road, Haidian District Beijing, 100191, P. R. China.

## Abstract

Mutant *RHO* is the most frequent genetic cause of autosomal dominant retinitis pigmentosa. Here, we developed an allele-specific gene editing therapeutic drug to selectively target the human T17M *RHO* mutant allele while leaving the wild-type *RHO* allele intact for the first time. We identified a *Staphylococcus aureus* Cas9 (SaCas9) guide RNA that was highly active and specific to the human T17M *RHO* allele. *In vitro* experiments using HEK293T cells and patient-specific induced pluripotent stem cells (iPSCs) demonstrated active nuclease activity and high specificity. Subretinal delivery of a single adeno-associated virus serotype 2/8 packaging SaCas9 and sgRNA to the retinas of the *RHO* humanized mice showed that this therapeutic drug targeted the mutant allele selectively, thereby downregulating the mutant *RHO* mRNA expression. Administration of this therapeutic drug resulted in a long-term (up to 11 months after treatment) improvement of retinal function and preservation of photoreceptors in the mutant humanized heterozygous mice. Our study demonstrated a dose-dependent therapeutic effect *in vivo.* Unwanted off-target effects were not observed at the whole-genome sequencing level. Our study provides strong support for the further development of this effective therapeutic drug to treat *RHO*-T17M associated autosomal dominant retinitis pigmentosa (adRP), also offers a generalizable framework for developing gene editing medicine. Furthermore, our success in retoring the vision loss in the suffering *RHO* humanized mice verifies the feasibility of allele-specific CRISPR/Cas9-based medicines for other autosomal dominant inherited retinal dystrophies.

## Introduction

Retinitis Pigmentosa (RP [MIM:268000]), a subtype of Inherited Retinal Dystrophies (IRDs), is one of the most common causes of blindness in industrialized countries, with an incidence of one in 1000 in North China (Xu et al., 2006). However, there are currently no reliable therapies available to cure or delay the disease progression of RP. In 2017, the US Food and Drug Administration approved Luxturna, an adeno-associated virus (AAV)-based gene therapy drug for *RPE65*-associated Leber’s Congenital Amaurosis (*RPE65*-LCA) (Russell et al., 2017, Smalley, 2017), which marked the start of gene therapy for IRDs (Lee et al., 2019).

Mutations in the human rhodopsin (*RHO*) gene are the most prominent genetic causes of autosomal dominant RP (adRP), accounting for about 30% of adRP cases(Iannaccone et al., 2006). However, different from the *RPE65*-LCA, which is caused by a loss-of-function mutation, *RHO*-adRP is mainly caused by a gain-of-function mutation(Li et al., 2018, Mendes et al., 2005), thereby restricting the application of gene replacement therapy.

Many studies have been conducted regarding the treatment technique for *RHO*-adRP, such as allele-independent gene ablation. This technique includes silencing the expression of both the mutant *RHO* allele and its wild-type (WT) counterpart and simultaneously adding an exogenous WT *RHO* gene beyond the regulation of the retina *in vivo*(Mitra et al., 2018, Cideciyan et al., 2018, Tsai et al., 2018). Although these methods are applicable to adRP associated with the *RHO* mutation, many risks remain with the disruption of the WT allele. Recently, allele-specific methods have been used extensively to treat adRP(Meng et al., 2020). With the aim of allele-specific *RHO* gene downregulation at the RNA level using antisense oligonucleotides (ASO), short hairpin RNA (shRNA), or RNA interference (RNAi) (Meng et al., 2020) can be achieved, some of which have been translated into clinical trials. However, repeated treatments, especially for sub-retinal injections, are bound to hamper therapeutic effectiveness. In comparison, allele-specific mutant DNA targeting approaches, specifically the with CRISPR/Cas9 system(Mali et al., 2013, Ran et al., 2015), which leads to permanent DNA modification, may be more promising. In fact, the CRISPR/Cas9 system has been used extensively to treat adRP caused by *RHO*-P23H, S334ter, P347S mutations, *etc.* This resulted in the delay in photoreceptor degeneration and rescue of retinal function in animal models(Latella et al., 2016, Li et al., 2018, Giannelli et al., 2018, Patrizi et al., 2021, Bakondi et al., 2016). However, these proof-of-concept studies are not suitable for direct translation into clinical practice. Therefore, there is currently no approved treatment for adRP.

The P23H mutation is the most common *RHO* mutation in North America, accounting for 10% of adRP cases due to a founder effect, while it has not been reported elsewhere, including Europe and Asia(Meng et al., 2020). Data gathered from our previous study(Liu et al., 2021b) showed that *RHO*-T17M is relatively common in China. In addition, several studies and data from the Human Gene Mutation Database (HGMD, http://www.hgmd.cf.ac.uk/ac/index.php) also prove that T17M(Sheffield et al., 1991, Krebs et al., 2010, Rakoczy et al., 2011) is one of the most common *RHO* mutations.

Previous *in vitro* and *in vivo* studies have shown that if the disease-causing allele has unique protospacer adjacent motif (PAM) sequences that are not present in the WT allele, or when the mutations are within the spacer region of the Cas9/single guide RNA (sgRNA) target site, the disease-causing allele may be discriminated from its WT counterpart. Therefore, the aim of selectively inactivating the mutant allele while leaving the WT ones functionally intact can be achieved(Burnight et al., 2017, Li et al., 2018, Giannelli et al., 2018, Patrizi et al., 2021). Non-homologous end joining (NHEJ) is the primary mechanism for double-strand breaks(DSBs) in terminally differentiated rod cells, whereas homologous recombination commonly occurs in mitotic cells such as induced pluripotent stem cells (iPSCs) or 293T (Chan et al., 2011). This difference in rod cells from mitotic cells emphasizes the significance and necessity of performing experiments *in vivo*, especially in humanized animal models. Unfortunately, allele-specific CRISPR/Cas9 experiments have not been performed in humanized animal models.

Hence, we developed an allele-specific gene editing therapy to selectively target the human T17M (c. 50C>T) *RHO* mutant allele while leaving the WT *RHO* allele intact for the first time. Our *in vitro* experiments using HEK293T cells and patient-specific induced pluripotent stem cells (iPSCs) demonstrated high nuclease specificity. Furthermore, an *RHO* humanized mouse model(Liu et al., 2021a) generated previously can simulate pathological changes in *RHO*-adRP. Using this mouse model, our *in vivo* experiments demonstrated specific inactivation of the mutant *RHO* allele and a delay in photoreceptor degeneration for up to 11 months after treatment. Our study demonstrated a dose-dependent therapeutic effect *in vivo*. Unwanted off-target effects were not observed. Thus, our study clearly provides strong evidence to support further development of this therapeutic drug for patients with *RHO-*adRP.

## Results

### Identification of allele-specific sgRNAs for the *RHO*-T17M allele

Our objective was to delete the mutant *RHO* allele using an allele-specific sgRNA against mutational sites with NNGRR PAMs when using SaCas9, thereby tethering the SaCas9 protein to the mutant but not the WT allele. The mutant allele, including the T17M mutation, did not generate unique PAM sequences. Allele-specific sgRNAs for T17M were designed to be within the spacer region of the Cas9/SgRNA target site. For T17M, three sgRNAs, 17-Sg1: 5’-CGTACCACACCCATCGCATTG-3’ (PAM: GAGAA), 17-Sg2:5’-TGCGTACCACACCCATCGCAT-3’ (PAM: TGGAG), and 17-Sg3: 5’-CTGCGTACCACACCCATCGCA-3’ (PAM: TTGGA) were designed to target the mutant allele (Fig.1A). *In vitro* digestion of either WT or mutant *RHO* sequence with SaCas9/SgRNA was carried out to assess the specificity profile of these 17-sgRNAs. 17-Sg1 and Sg2 appeared to preferentially cut the mutant sequence, but not the WT sequence, whereas 17-Sg3 cut both the WT and mutant sequences and appeared to cut the mutant sequence more than the WT sequence (Fig. 1B).

**Fig. 1.**
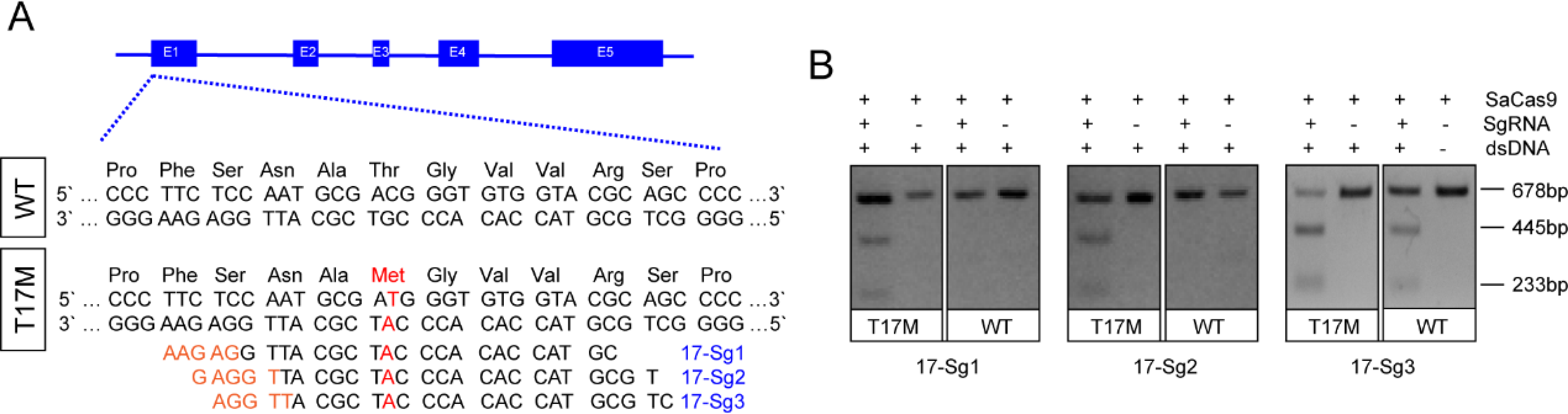
Schematic representation of allele-specific sgRNA design for *RHO*-T17M (A) and confirmation of the specificity achieved using SaCas9 protein complexed with a sgRNA to target the mutant *RHO* sequence and its corresponding wildtype sequence *in vitro* (B), the full-length amplicon was 678bp, the two truncated amplicons were 445 bp and 233 bp, respectively. The mutation c.50C>T (p.T17M) is indicated in red. Protospacer adjacent motif sequences were marked in orange. Exons were indicated by closed boxes.

### *In vitro* knockdown of *RHO*-T17M gene expression by allele-specific gene editing in 293T cells

Due to the unavailability of cell lines expressing human *RHO*, we created 293T cell pool stably expressing *RHO*-WT or *RHO*-T17M using lentivirus FUGW-*RHO*-cDNA (Fig.2A). Subsequently, 293T cells expressing *RHO*-WT (RHO*wt* cells) and RHO17 cells were transfected with pX601-EFS-SaCas9-U6-17-Sg1 (17-Sg1) or Sg2 plasmid (Fig.2A). Notably, RHO17 cells that were not transfected assayed with T7E1 nuclease only resulted in full-length amplicon (760 bp, Fig.2B). Similarly, T7E1 assay for RHO*wt* cells transfected with 17-Sg1 or 17-Sg2 plasmid, resulted in full-length amplicon (Fig.2B). In contrast, the T7E1 assay for RHO17 cells transfected with 17-Sg1 or Sg2 plasmid demonstrated that 17-Sg1 and Sg2 were able to specifically cut the mutant sequence (Fig.2B). Subsequent TA cloning (59 clones) were chosen for each sgRNA) and Sanger sequencing were used to measure the frequency of insertion or deletion mutations (indels) . Indels were not detected in WT samples. However, indels were found at frequencies of 11.86% (7/59) and 25.42% (15/59) in samples of SaCas9/17-Sg1 and SaCas9/17-Sg2, respectively (Fig. 2C, Supplementary Table S1). Moreover, we evaluated rhodopsin protein production through Western Blotting (WB). Rhodopsin expression in RHO17 cells transfected with 17-Sg1 or Sg2 plasmid were strongly reduced compared to the counterpart of RHO*wt* cells with 17-Sg1 or 17-Sg2 plasmid (Fig. 2D) .

**Fig. 2.**
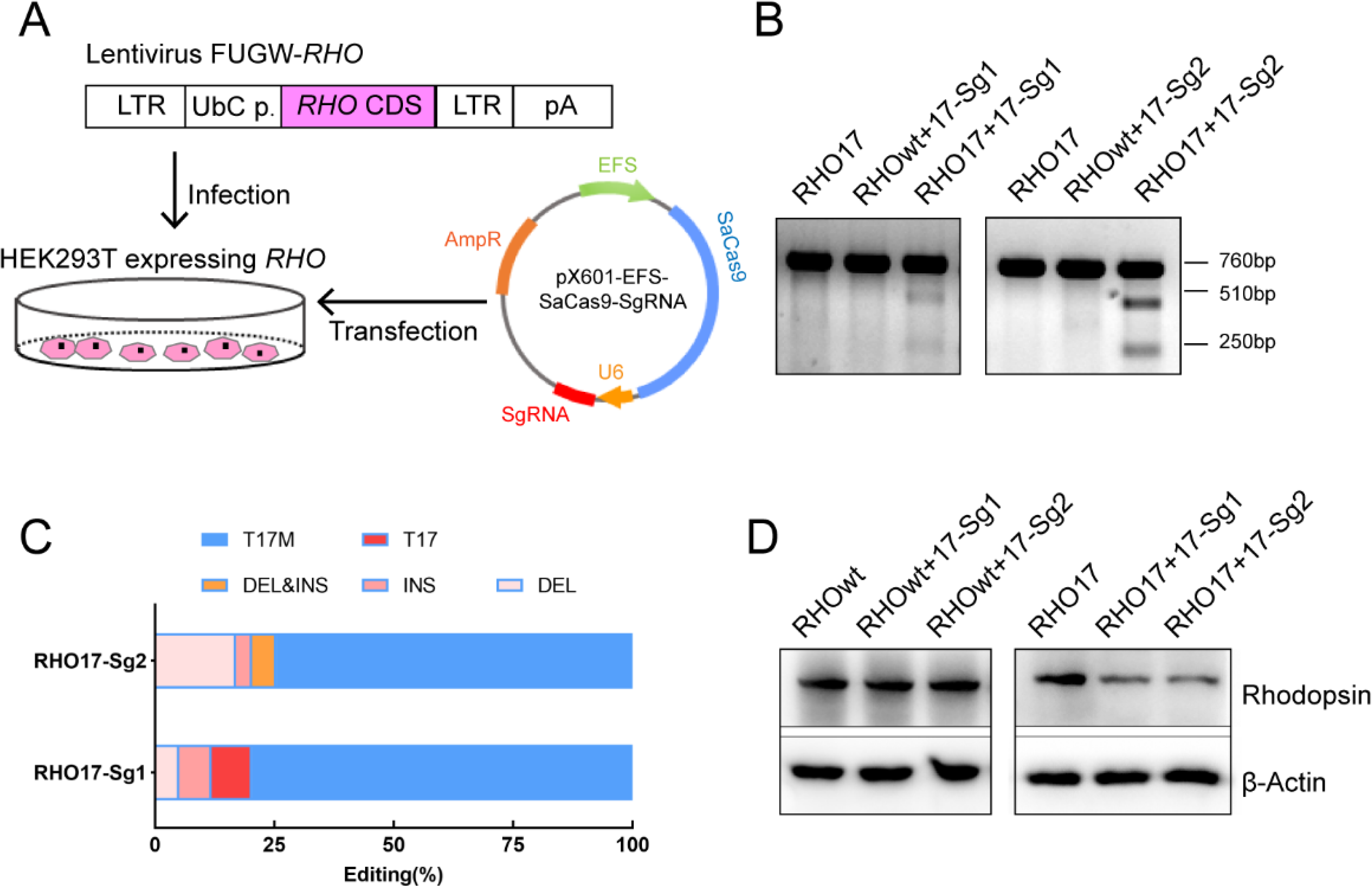
In v***i***tro **knock-down of human *RHO-*T17M expression.** (A) Schematic view of construction of 293T stably expressing human *RHO* protein and transfection of pX601-EFS-SaCas9-U6-sgRNA (SaCas9/SgRNA) plasmid. (B) T7E1 assay indicated that SaCas9/17-Sg1 and SaCas9/17-Sg2 were appeared to cut the mutant sequence specifically, the full-length amplicon was 760bp, the two truncated amplicons were 510 bp and 250 bp, respectively. (C) The cutting efficacy of two sgRNAs with SaCas9 determined by TA & Sanger sequencing in 293T cells. (D) *Rhodopsin* expression reduction was determined by western blotting in 293T cells transfected with *RHO*-T17M-pEGFPN1(RHO17 cells) transfected with 17-Sg1 and Sg2 plasmid, comparing to the RHO*wt* cells with 17-Sg1 and Sg2 plasmid.

To further analyze the effect of allele-specific knockdown of the *RHO*-T17M, we generated 293T cells that expressed SaCas9/17-Sg1 or SaCas9/17-Sg2 using lentivirus SaCRISPR_SgRNA-GFP. These 293T cells were then transfected with *RHO*-pmCherryN1 plasmids with or without the T17M mutation (Supplementary Fig. S1A). Seventy-two hours post transfection, fluorescence intensity was examined, and the mCherry/GFP fluorescence intensity ratio (mCherry/GFP ratio) was assessed. There was no significant difference in the mCherry/GFP ratio between 293T cells expressing only SaCas9 (Lenti-CTRL cells) transfected with *RHO*-WT-pmCherryN1 (RHO*wt*) plasmid, and 17-Sg1 or 17-Sg2 cells with RHO*wt* plasmid (Supplementary Fig. S1B-1C). The mCherry/GFP ratio in 17-Sg1 and 17-Sg2 cells transfected with the RHO17 plasmid was significantly reduced compared to that of Lenti-CTRL cells transfected with the RHO17 plasmid (Supplementary Fig. S1D-1E). These results evidently indicated that SaCas9/17-Sg1 and Sg2 enabled discrimination between WT and mutant alleles, targeting the mutant allele exclusively. The 17-Sg2 had higher cutting effiency than 17-Sg1 and was chosen for further *in vitro* and *in vivo* experiments.

### Allele-Specific gene editing using CRISPR/Cas9 in patient-derived iPSCs

Urine cells (UCs, Fig. 3A) from Patient 1 (P1) diagnosed with RP(Liu et al., 2021b) caused by *RHO*-T17M mutation were obtained and reprogrammed to iPSCs to further determine the specificity and cutting efficiency of SaCas9/17-Sg2 in patient cells. Patient-derived iPSCs manifested a typical cobblestone-like morphology (Fig. 3B). Sanger sequencing confirmed that patient P1 carried the heterozygous variant c.50C>T (p.T17M) in *RHO* (Fig. 3C-3D). Immunofluorescence (IF) staining showed that their iPSCs expressed human embryonic stem cell-specific surface antigens(International Stem Cell et al., 2007) including NANOG, OCT4, TRA-1-60, and SSEA-4 proteins (Fig. 3E-H). Chromosomal content analysis revealed a normal 46, XY karyotype (Fig. 3I). P1 iPSCs (P1 iPS) and iPSCs from a normal donor (NoR iPS) were transfected with the AAV-EFS-SaCas9-U6-17-Sg2-p2a-Puro (17-Sg2-Puro) plasmid (Fig. 3J). Genomic DNA from these iPS cells was subjected to T7E1 assay. The results indicated that 17-Sg2 appeared to specifically cut the mutant sequence (Fig. 3K). Hi-Tom sequencing revealed that the cutting efficiency of 17-Sg2 was 3.94% (Fig. 3L and Supplementary Fig. S2A). To further confirm this allele-specific cutting, colonies from P1 iPS were picked and transfected with the 17-Sg2-Puro plasmid. Similarly, a cutting efficiency of 2.47% was dected (Supplementary Fig. S2B). A 3bp-deletion (c.44_46del3bp) in the *RHO*-T17M allele was found, whereas no indels were found in the *RHO*-WT allele. Moreover, 17-Sg2 did not facilitate SaCas9-mediated cleavage of the WT allele (Fig. 3K), and indels were not found (Fig. 3M) in transfected NoR iPS. These results firmly indicate that the mutant allele-specific gRNA 17-Sg2 targets the mutant allele specifically in patient derived iPSCs.

**Fig. 3.**
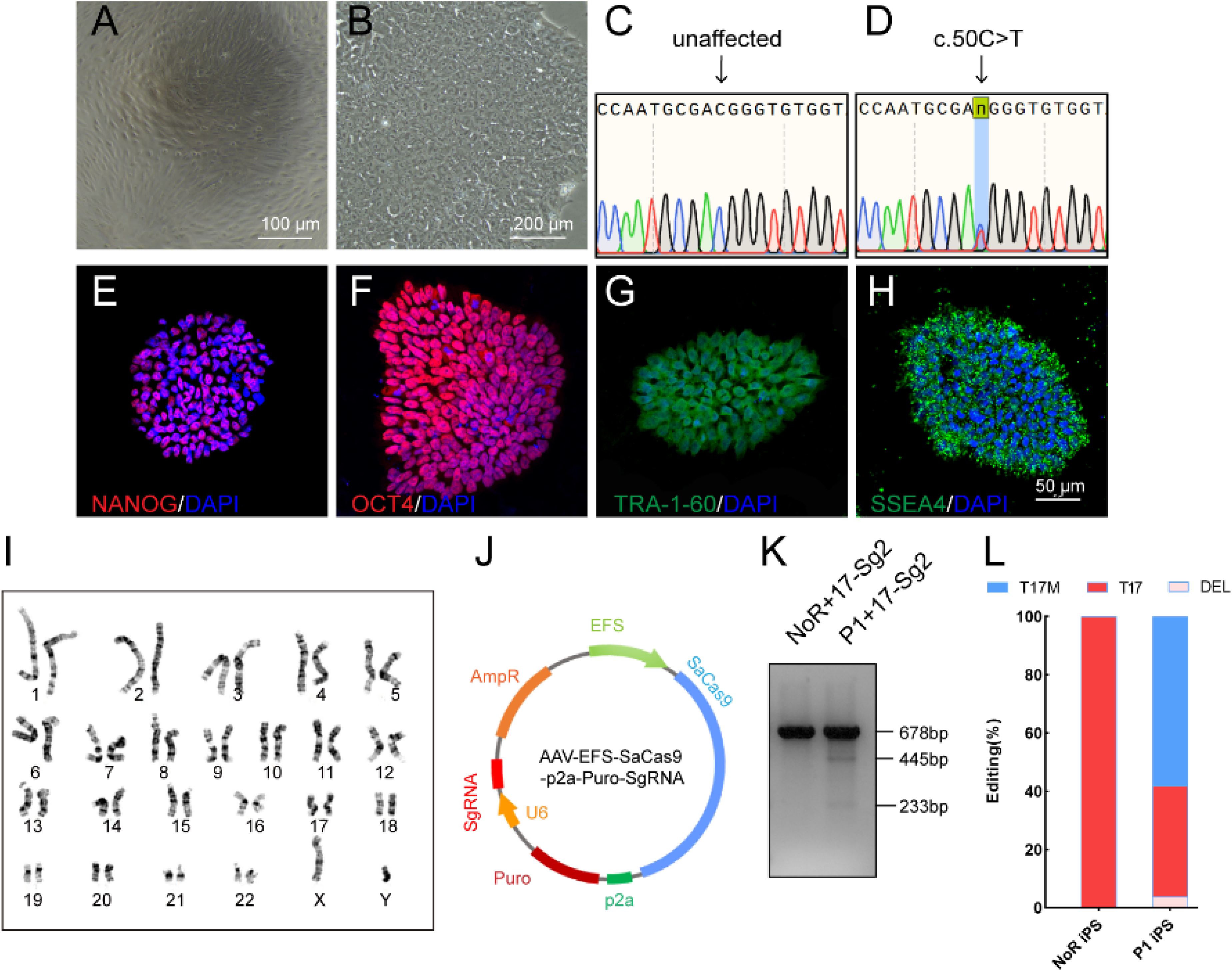
Determination of specificity and cutting efficiency of SaCas9 and sgRNA in induced pluripotent stem cells (iPSCs) from patients’ urine cells (UCs). (A)Photograph of UCs. Scale bar=100 μm. (B) Photograph of iPSCs. Scale bar = 200 μm. (C-D) Sanger sequencing results showed that P1 iPS had the heterozygous c.50C>T mutation. IF staining showed the iPSCs expressed human embryonic stem cell-specific surface antigens including NANOG (E), OCT4 (F), TRA-1-60 (G) and SSEA-4 (H) protein. (I) Chromosomal content analysis revealed a normal 46, XY karyotype of patient P1. (J) Schematic view of AAV-EFS-SaCas9-U6-sgRNA-p2a-Puro plasmid. (K) T7E1 assay indicated that SaCas9/17-Sg2 was appeared to cut the mutant sequence specifically. (L) The cutting efficacy of SaCas9/17-Sg2 is determined by TA & Sanger sequencing in iPSCs.

### Specific editing of the mutant *RHO* allele in humanized mice

After confirming the ability of 17-Sg2 to target the *RHO*-T17M allele specifically in both 293T and patient derived iPSCs, we aimed to develop gene therapy drug using 17-Sg2 and SaCas9. We chose AAV2/8 as the vector for the drug development because of its superior infectivity to photoreceptor cells. As the first step, we tested if the EFS promoter was an appropriate promoter to drive strong gene expression in the mouse retina using GFP as a reporter. We constructed AAV2/8-EFS-EGFP (AAV-EGFP) and subretinally injected it into both eyes of the *Rho^wt/wt^* mice at either high (3×10^9^ vg per eye, 1 μL) or low (1×10^9^ vg per eye, 1 μL) dose (Supplementary Table S2, Fig. 4A). Injection blebs were imaged immediately following injection (Fig. 4B). One month post injection (1 mpi), the neural retinas of the mice were dissected and GFP positive (GFP+) and negative (GFP-) areas were analyzed, as illustrated in Fig. 4C. IF analysis indicated that GFP was expressed in the OS/IS and ONL of photoreceptor cells at 1 mpi and 11 mpi (Fig. 4D). These data indicated that the EFS promoter drived sufficient gene expression in photoreceptors, meeting the requirement for therapeutic drug development. The to evaluated whether the EFS promoter could similarly express SaCas9 and in the mouse retina, we construced an expression vector AAV2/8-EFS-SaCas9-U6-17-Sg2 (AAV-SaCas9/17-Sg2) and produced AAV viruses. To recognizing the subretinal injection sites and considering the limited cargo capacity of AAV, AAV-EGFP and AAV2/8-EFS -SaCas9/17-Sg2 (1:1 mixture) were co-delivered into both eyes of the *Rho^wt/hum^* mice at different doses, each at 5×10^8^ (1×10^9^ vg per eye totally, 1 μL), 1×10^9^ (2×10^9^ vg per eye totally, 1 μL) and 3×10^9^ (6×10^9^ vg per eye totally, 2 μL) dose (Fig. 4A, Supplementary Table S2). As shown in Fig 4E and 4F, qPCR analysis revealed that 17-Sg2 and SaCas9 were expressed in all injected retinas at 3 mpi. WB analysis indicated that SaCas9 was present in injected retinas at 3 mpi (Fig. 4G) at doses of 1×10^9^ and 3 × 10^9^.

**Fig. 4.**
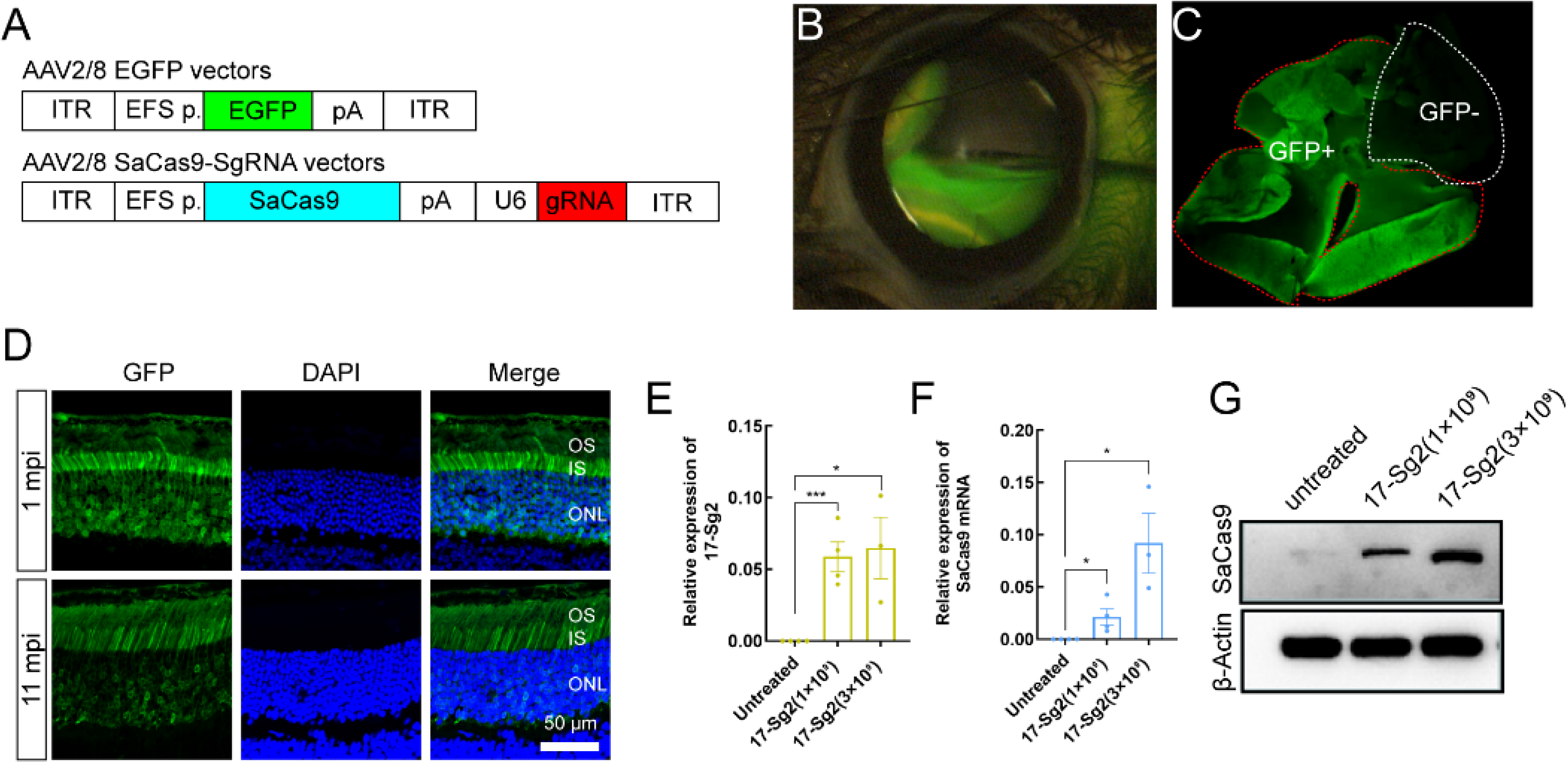
AAV2/8-mediated SaCas9 and sgRNA expression in mouse retinas. (A) (Top) AAV2/8 vectors expressing EGFP under the control of EFS promoter; (Bottom) AAV2/8 vectors expressing SaCas9 under the control of EFS promoter and sgRNA under the control of U6 promoter were schematized. (B) Fluorescein shows AAV vectors distribution following subretinal injection immediately. (C) Infected areas treated with AAV2/8-EFS-EGFP vector at dose of 1×10^9^ vg per eye (1×10^9^ dose) were labelled by GFP expression (GFP+ area). ONL = outer nuclear layer; OS = outer segment; IS = inner segment. Scale bar = 50 μm. (D) IF staining indicates GFP expresses in mouse retinas at 1m and 11 mpi treated with AAV2/8-EFS-EGFP vector at 3×10^9^ dose. (E) The quantitative PCR analysis indicated the expression of SaCas9, 17-Sg2 and GFP in *Rho^wt/hum^* retinas at 3 mpi treated with AAV2/8-EFS-SaCas9-U6-17-Sg2 and AAV2/8-EFS-EGFP vector (1:1 mixture) at different doses. (F) Determination of SaCas9 expression by western blotting at 3 mpi at 1×10^9^ and 3×10^9^ dose. (Left) Mouse #h, i and j (*Rho^wt/hum^*) were injected with AAV2/8-EFS-SaCas9-U6-17-Sg2 and AAV2/8-EFS-EGFP vector (1:1 mixture) at 1×10^9^ dose; (Right) Mouse #k, l, m and n (*Rho^wt/hum^*) were injected with AAV2/8-EFS-SaCas9-U6-17-Sg2 and AAV2/8-EFS-EGFP vector (1:1 mixture) at 3×10^9^ dose.

Previously, we generated *RHO* humanized mouse models (Liu et al., 2021a) using CRISPR/Cas9 and donor plasmids by replacing the mouse *Rho* gene with the human counterpart (Supplementary Fig. S3A-S3B). *Rho^wt/hum^* mice was heterozygotes with a mouse *Rho*-WT allele and human *RHO*-WT allele, and *Rho^hum/hum^* mice were homozygotes with human *RHO*-WT allele. Mut-*Rho^wt/hum^* mice were heterozygotes with a mouse *Rho*-WT allele and human *RHO*-5m allele, Mut-*Rho^hum/hum^* mice were homozygotes with *RHO*-5m allele.To demonstrate the ability of the allele-specific SaCas9/SgRNA system to edit the human *RHO*-T17M allele *in vivo*, the Mut-*Rho^wt/hum^* mouse model was used to test cutting efficiency and *Rho^wt/hum^* to evaluate gene editing specificity (Supplementary Table S2). AAV-EGFP and AAV-SaCas9/17-Sg2 (1:1 mixture) were co-delivered into both eyes of Mut-*Rho^wt/hum^* and *Rho^wt/hum^* mice, each at 5×10^8^, 1×10^9^ and 3×10^9^ dose. GFP+ and GFP-areas of injected retinas or whole retinas (indicating GFP+ areas could be observed in all retinas) were harvested at 3 mpi. T7E1 assay showed that SaCas9/17-Sg2 was able to cut the mutant sequence specifically (Fig. 5A, Supplementary Fig. S4A-S4B). Truncated cleaved products were detected in Mut-*Rho^wt/hum^* mice retinas (mouse #M1∼M10) but not in *Rho^wt/hum^* mice retinas (mouse #C1∼C8) at three different doses, but no truncated cleaved products were detected in two mouse retinas (5×10^8^ dose, data not shown). The average efficiencies calculated using TA and Sanger sequencing were 3.33% (GFP+, *n* = 1), 13.99% (GFP+, *n* = 4),16.59% (GFP+, *n* = 5) in the 5×10^8^, 1×10^9^, and 3×10^9^ dose groups, respectively (Supplementary Table S3). Hi-Tom sequencing indicated the average cutting efficiencies of 17-Sg2 were 6.37% (GFP+, *n* = 1) and 1.60% (GFP–, *n* = 1), 11.27% (GFP+, *n* = 4) and 6.10% (GFP–, *n* = 4), 32.58% (GFP+, *n* = 5) and 11.04% (GFP–, *n* = 5) at 5×10^8^, 1×10^9^ and 3×10^9^ dose, respectively (Fig. 5B). PCR-based NGS indicated the average efficiencies were 6.85% (GFP+, *n* = 1), 14.20% (GFP+, *n* = 4), 16.25% (GFP+, *n* = 5) in the 5×10^8^, 1×10^9^, and 3×10^9^ dose groups, respectively (Fig. 5C-5D). Therefore, the cutting efficiency showed a dose response. We observed that 89.53% (77/86, TA& Sanger sequencing) of the indels were frameshifts (Supplementary Table S3), which were expected to create a premature termination codon (PTC) and thereby resulted in the nonsense-mediated decay (NMD) of the edited *RHO*-5m mRNA (https://www.genecascade.org/MutationTaster2021). Among these, c.46_46delG (p. A16Rfs*32), the most common deletion, accounted for 20.93% (18/86) of all indels. We found that the translation of majority of the edited *RHO*-5m protein could be stopped due to PTC (Fig. 5E).

**Fig. 5.**
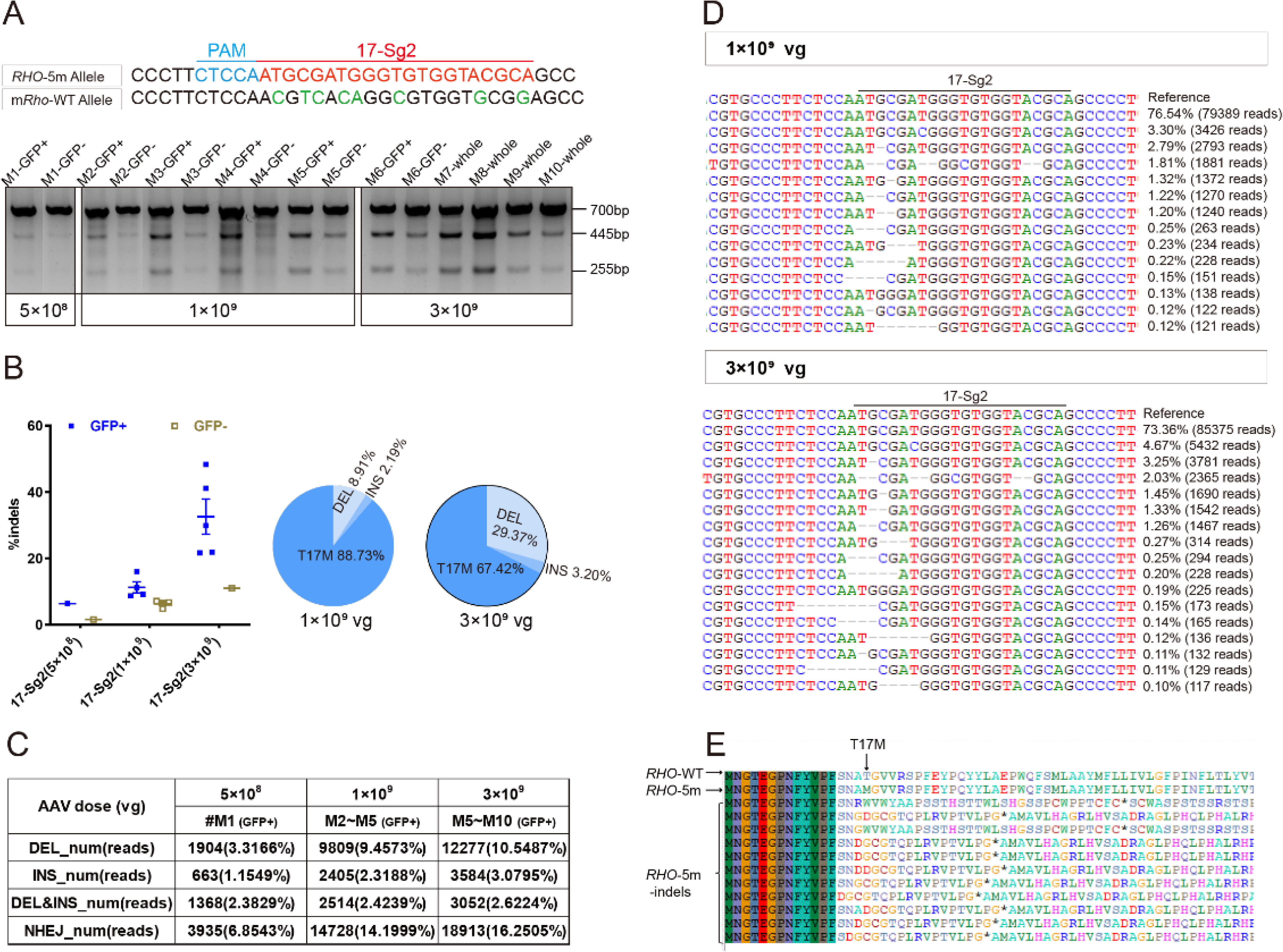
CRISPR/Cas9 targeting of c.50C>T *RHO* dominant variant encoding p.T17M in Mut-*Rho^wt/hum^* mouse retinas. (A) T7E1 assay was performed on GFP+ and GFP-retinas to detect gene editing activity of AAV2/8-EFS-SaCas9-U6-17-Sg2 (AAV2/8-SaCas9/17-Sg2) in Mut-*Rho^wt/hum^* retinas at 5×10^8^, 1×10^9^ and 3×10^9^ doses, the full-length amplicon was 700bp, the two truncated amplicons were 445 bp and 255 bp, respectively; 17-Sg2 was marked in red; Protospacer adjacent motif sequence was marked in light blue and mismatches in mouse *Rho* sequence comparing to 17-Sg2 were indicated in green. Hi-Tom sequencing (B) and PCR-based Next-generation sequencing (C) were used to detect the gene editing efficiency and types of indels in AAV2/8-SaCas9/17-Sg2-treated Mut-*Rho^wt/hum^* retinas at 5×10^8^, 1×10^9^ and 3×10^9^ doses. (D) Graphic representation of indels scored in the target site of Mut-*Rho^wt/hum^* retinas treated with AAV2/8-SaCas9/17-Sg2 at 1×10^9^ and 3×10^9^ doses. (E) Analysis of the amino acid sequence of the edited *RHO*-5m allele, * indicated the premature termination codon.

To determine the off-target activity of 17-Sg2 in the mouse *Rho* allele (8 mismatches between 17-Sg2 and the corresponding mouse *Rho* sequence (Supplementary Fig. S4C), T7E1 assay was performed and the data clearly showed that SaCas9/17-Sg2 did not cut the mouse *Rho* sequence (Supplementary Fig. S4C). To provide additional support evidence, AAV-EGFP and AAV-SaCas9/17-Sg2 (1:1 mixture) were similarly co-delivered into the eyes of Mut-*Rho^hum/hum^* mice at 1×10^9^ and 3×10^9^ dose (Supplementary Table S2). However, the Mut-*Rho^hum/hum^* mice suffered from extremely rapid degeneration of photoreceptor cells and defect formation of OS which was caused by retaining the *RHO*-5m proteins in RIS and cell bodies (Supplementary Fig. S4D). Moreover, there was no cutting activity detected in the T7E1 assay (Supplementary Fig. S4E). This result might indicate that infected photoreceptor cells all died at 3 mpi (the age of the mice was approximately P4.5m), photoreceptor cells were terminally differentiated neurons without regeneration capacity, and their existence was essential for the application of gene therapies.

These results from humanized mouse models obiviously demonstrated that 17-Sg2 specifically target the *RHO*-T17M allele and might restore the retinal function and vision loss in these mice models.

### Decreased expression level of the *RH*O-5m mRNA in AAV2/8-SaCas9/17-Sg2-treated *Rho^hum/m-hum^* retinas

To verify whether the indels created by the activity of SaCas9/17-Sg2 in Mut-*Rho^wt/hum^* mice could lead to attenuation or ablation of the *RHO*-5m allele, we cloned the three most frequent deletions—c.46_46delG (p. A16Rfs*32), c.45_46delTG (p. A16Dfs*17), and c.43_46delAATG (p. N15Rfs*32) and an insertion mutation (c.46_47ins49bp; p. A16Gfs*34) in the edited allele (Fig. 6A) — overexpressed them as well as *RHO*-WT and *RHO*-T17M in 293T cells. After one week of transfection, IF staining using 4D2 antibody (antibody recognizing N-terminal of rhodopsin protein, red) showed that the *RHO*-WT protein was localized on the plasma membrane, while the *RHO*-T17M protein was retained in the cytoplasm in 293T cells (Fig. 6B). The edited *RHO*-T17M protein of these four indels were undetectable using the 4D2 antibody (Fig. 6B). Similar results were obtained for the *RHO*-5m variant plasmid. Twenty-four hours after transfection, the number of GFP+ cells per 5000 μm^2^ in 293T cells transfected with different variant plasmids was statistically lower than that in 293T cells with the *RHO*-5m-PEGFPN1 plasmid (Supplementary Fig. S5A-5B). One week after transfection, IF staining using the 4D2 antibody showed that the *RHO*-5m protein was retained in the cytoplasm of 293T cells, while the *RHO*-5m protein of these four indels were undetectable using the 4D2 antibody (Fig. 6C). GFP+ cells per random sight were counted, and the percentage of GFP+ cells expressing rhodopsin in different variant plasmid groups was calculated (Fig. 6D), the number of GFP+ cells in the different variant plasmid groups was statistically lower than that in the *RHO*-5m-PEGFPN1 plasmid group, and rhodopsin was rarely expressed in 293T cells transfected with different variant plasmids (Fig. 6E-6F). Taken together, the indels that occurred in the *RHO*-5m allele led to frameshifts, consistent with the results by *in silico* analysis. The *in vitro* results also indicated that although five variants in the mutant *RHO* allele in Mut-*Rho^wt/hum^* mouse, indels created by gene editing around the T17M mutation (variant closest to the N terminal of rhodopsin) could lead to the ablation of the *RHO*-5m mRNA. The existence of the other four variants in the *RHO* coding region would not interrupt the allele specific gene editing for T17M. These data indicated that Mut-*Rho^wt/hum^* mouse could be a practical model that can be used to evaluate phenotype rescues after treatment.

**Fig. 6.**
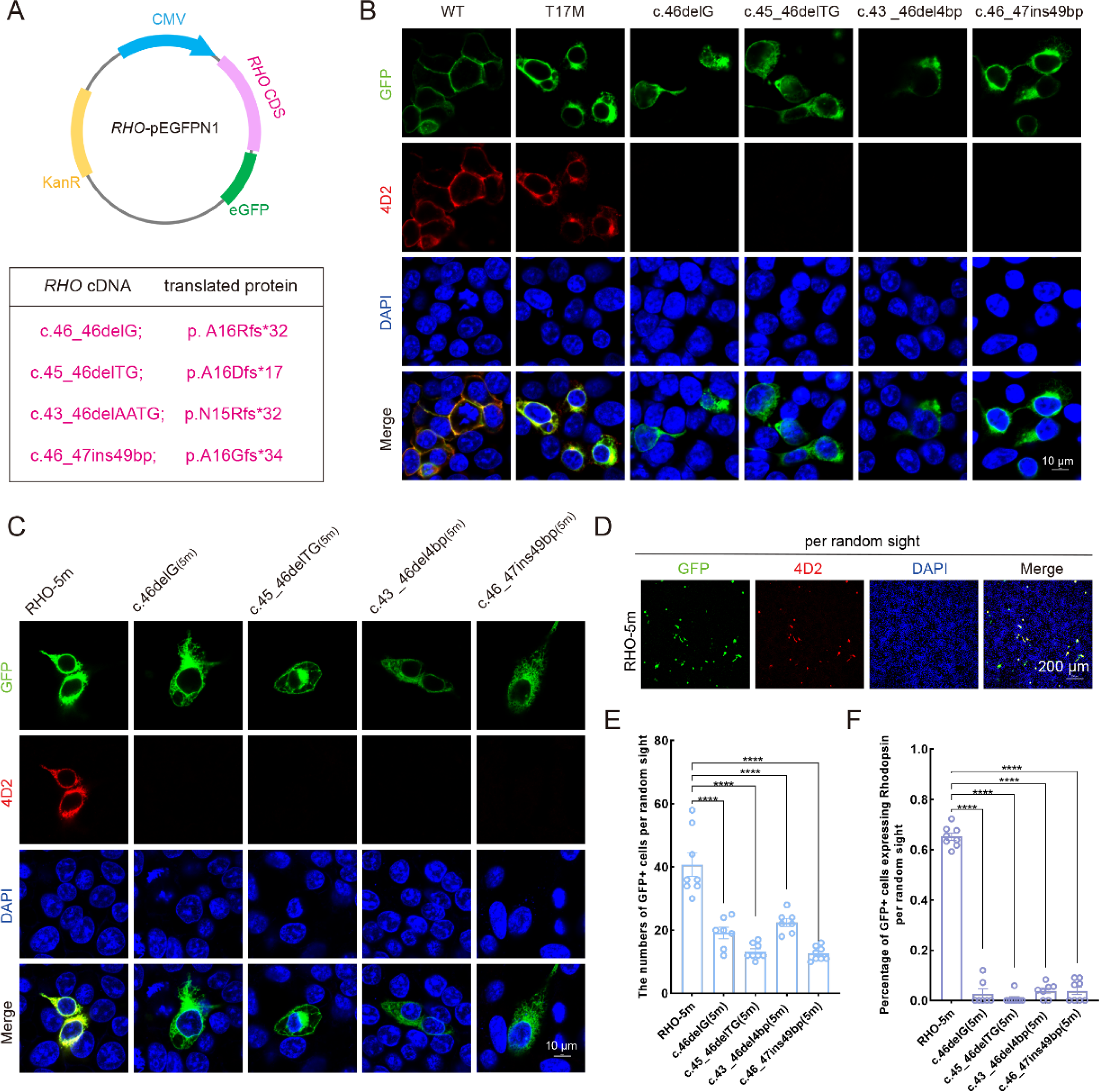
Expression of the mutant human *RHO* allele after gene editing with SaCas9/17-Sg2 *in vitro*. (A) Schematic view of the different human *RHO* gene variants created by gene editing. (Top) map of the pEGFPN1 vector used to overexpress these variants; (Bottom) the description of variants at DNA and protein level. (B) Colocalization of GFP and rhodopsin (4D2, red) in HEK293T cells transfected with pEGFPN1 vector carrying *RHO*-WT, *RHO*-T17M and four edited *RHO*-T17M variants, one week after transfection. Scale bar = 10 μm. (C) Colocalization of GFP and rhodopsin (4D2, red) in 293T cells transfected with pEGFPN1 vector carrying *RHO*-5m and four edited *RHO*-5m variants, one week after transfection, Scale bar = 10μm. (D-F) The number of GFP+ cells and percentage of GFP+ cells expressing rhodopsin per random sight, Scale bar = 200 μm. Nuclei were stained blue by DAPI.

We then determined whether the indels created by SaCas9/17-Sg2 at the DNA level could result in decreased mRNA levels of the *RHO*-5m allele. AAV-EGFP and AAV-SaCas9/17-Sg2 (1:1 mixture) were co-delivered into both eyes of the *Rho^hum/m-hum^* mice, each at 1×10^9^ and 3×10^9^ doses (Supplementary Table S2). *Rho^hum/m-hum^* mouse, a humanized mouse model carried a human *RHO*-WT allele and *RHO*-5m allele. Truncated cleaved products treated with T7E1 nuclease were observed in treated *Rho^hum/m-hum^* retinas (Fig. 7A), the average cutting efficiencies were 10.06% (GFP+, *n* = 4 at 1×10^9^ dose) and 12.09% (GFP+, *n* = 4 at 3×10^9^ dose) revealed by Hi-Tom sequencing (Fig. 7B). Meanwhile, we employed Hi-Tom sequencing, TA and Sanger sequencing of RT-PCR products to compare the amount of mRNA transcripts of *RHO*-WT and *RHO*-5m in *Rho^hum/m-hum^* retinas at 3 mpi. At 1×10^9^ dose, Hi-Tom sequencing indicated that the average percentage of WT transcripts was higher in treated retinas than that in untreated retinas (53.31% *vs.* 48.51%), and the average mutant transcript was lower in treated retinas than that in untreated retinas (46.69% *vs.* 51.49%) (*p* = 0.05; Fig. 7C). A statistically higher percentage of WT transcripts and a lower percentage of mutant transcripts in treated retinas than the control (untreated retinas) were detected by TA and Sanger sequencing (*p*<0.05, Fig. 7D). At 3×10^9^ dose, Hi-Tom sequencing, TA and Sanger sequencing indicated that the percentage of WT transcripts was statistically higher in treated retinas than that in untreated retinas (*p*<0.005 and *p*<0.05, respectively), and the mutant transcript was statistically lower in treated retinas than that in untreated retinas (*p*<0.005 and *p*<0.05, respectively, Fig. 7C-7D). The qPCR analysis indicated that the relative *RHO* mRNA expression levels were significantly lower in the treated retinas than that in untreated retinas (*p*<0.05), at both 1×10^9^ and 3×10^9^ doses (Fig. 7E).

**Fig. 7.**
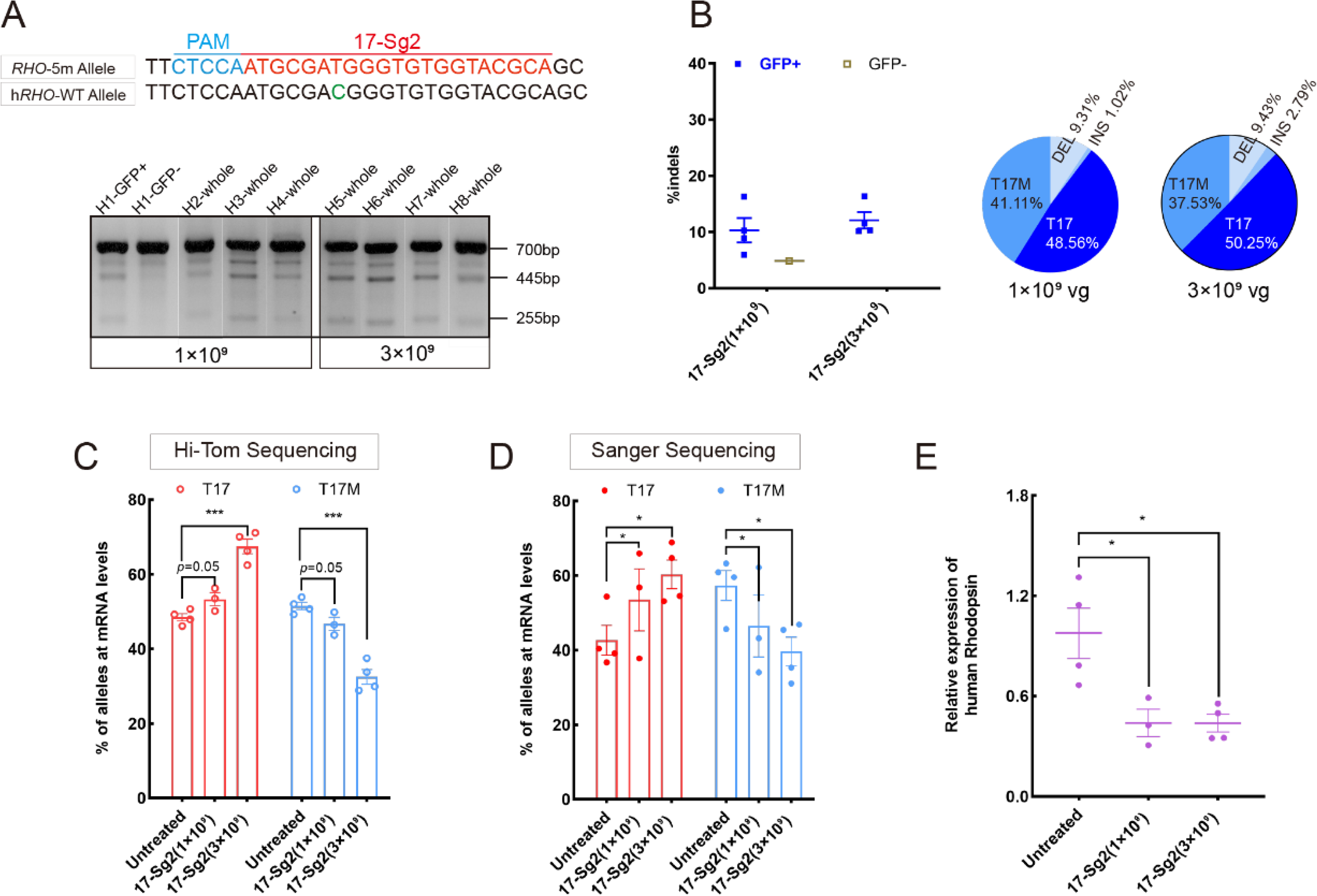
Expression of the *RHO*-5m allele after gene editing with AAV-based SaCas9/17-Sg2 in *Rho^hum/m-hum^* retinas. (A) T7E1 assay was performed on GFP+ and GFP-retinas to detect cutting activity of AAV2/8-EFS-SaCas9-U6-17-Sg2 (AAV2/8-SaCas9/17-Sg2) in *Rho^hum/m-hum^* retinas at 3 mpi, the full-length amplicon was 700bp, the two truncated amplicons were 445 bp and 255 bp, respectively; 17-Sg2 was marked in red; Protospacer adjacent motif sequence was marked in light blue and mismatches in wild type human *RHO* sequence comparing to 17-Sg2 were indicated in green. (B) Hi-Tom sequencing was used to detect the gene editing efficiency and types of indels in AAV2/8-SaCas9/17-Sg2-treated *Rho^hum/m-hum^* retinas at 1×10^9^ and 3×10^9^ doses. (E) Relative RNA level of WT and the *RHO*-5m alleles in SaCas9/17-Sg2-treated retinas (*n*=3 at 1×10^9^ dose, *n*=4 at 3×10^9^ dose) versus age-matched untreated retinas(*n*=4) determined by Hi-Tom Sequencing(E) and TA & Sanger Sequencing(F). Relative mRNA levels of human *RHO* in SaCas9/17-Sg2-treated retinas (*n*=3 at 1×10^9^ dose, *n*=4 at 3×10^9^ dose) versus age-matched untreated retinas(*n*=4) determined by qPCR analysis. Error bars show SEM, and the significance was calculated using two-tailed unpaired t-test, \**p*<0.05; ** *p*<0.01; *** *p*<0.005.

### Retinal functions improvement and photoreceptor preservation following the mutant *RHO* allele disruption

Scotopic electroretinogram (ERG) responses in Mut-*Rho^wt/hum^* mice were more affected than photopic responses (Supplementary Fig. S3C), indicating that rod dysfunction was more severe than cone dysfunction and deteriorated earlier. The ERG results were consistent with the HE and IF results, which showed that Mut-*Rho^wt/hum^* mice lost photoreceptors slowly, and the outer and inner segment (OS/IS) length of photoreceptor cells became shorter, with increasing age (Supplementary Fig. S3D-S3F), indicating the pattern of retinal degeneration caused by mutant human rhodopsin was a typical rod-cone decay. Thereby Mut-*Rho^wt/hum^* mice were affected with *RHO*-5m-adRP (Liu et al., 2021a).

To assess whether the retinal functions in treated mice improved, AAV-EGFP and AAV-SaCas9/17-Sg2 (1:1 mixture) or AAV-EGFP and AAV-SaCas9/CTRL (1:1 mixture) were co-delivered into both eyes of Mut-*Rho^wt/hum^* mice, each at 1×10^9^ and 3×10^9^ dose (Supplementary Table S2). 3 mpi, at doses of 1×10^9^ or 3×10^9^, ERG analysis showed a significant improvement in scotopic a- and b-waves in retinas treated with SaCas9/17-Sg2 compared to their corresponding controls at different stimulus intensities (Fig. 8A-8D). Similarly, 11 mpi, at 3×10^9^ dose, ERG analysis also indicated a significant improvement in scotopic a- and b-waves in retinas treated with SaCas9/17-Sg2 compared to their corresponding controls at several stimulus intensities (Fig. 8E-8F).

**Fig. 8.**
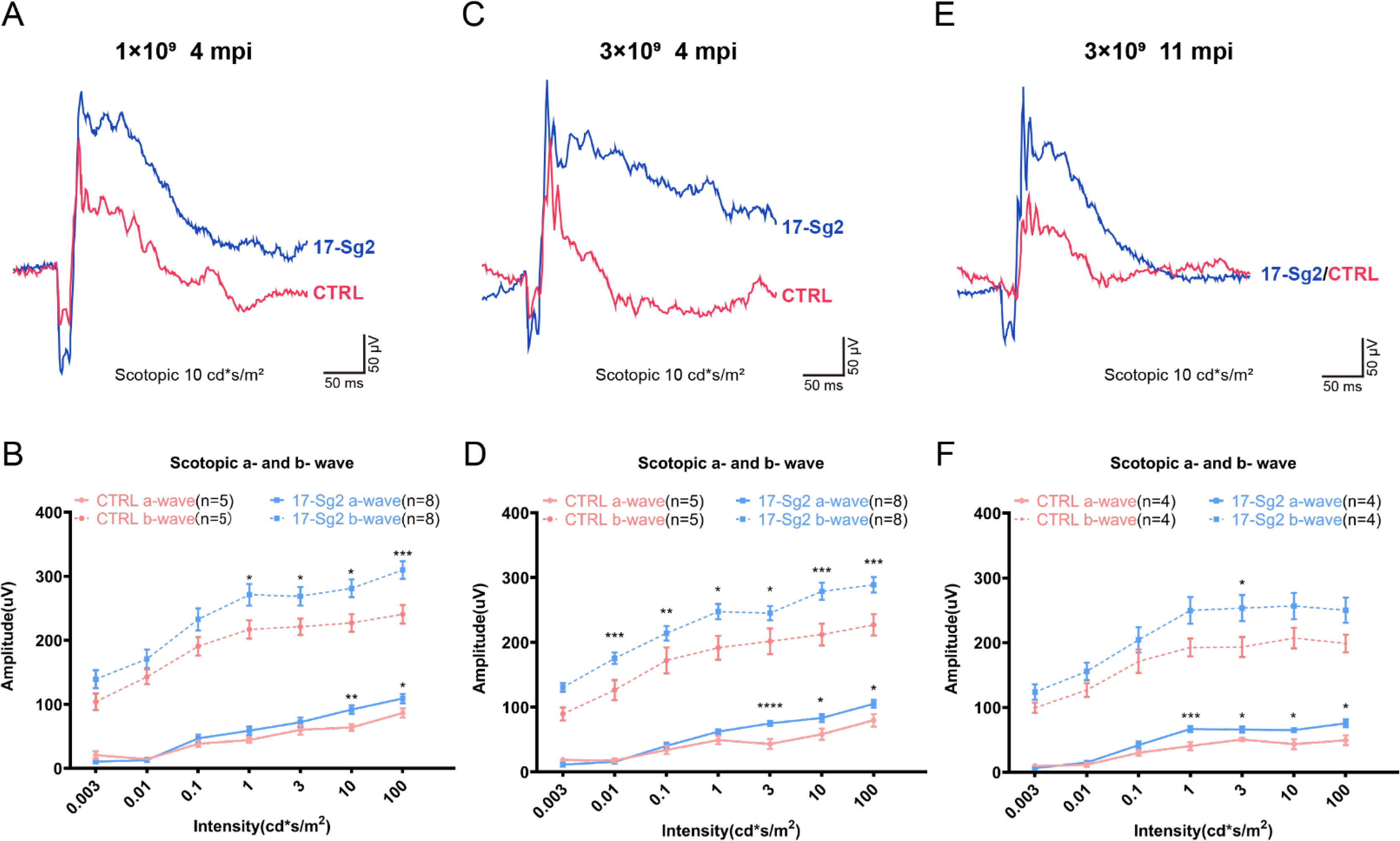
Significant improvement of retinal functions with AAV-based SaCas9/17-Sg2 in Mut-*Rho^wt/hum^* mice determined by full-field electroretinogram (ERG). Representative scotopic ERG (at the flash intensity of 10 cd*s/m^2^) waveforms of Mut-*Rho^wt/hum^* mice treated with AAV2/8-EFS-SaCas9-U6-17-Sg2 (AAV-SaCas9/17-Sg2, blue) and AAV-SaCas9/CTRL (red) at 1×10^9^ dose (3 mpi, A), 3×10^9^ dose (3 mpi, C) and at 3×10^9^ dose (11 mpi, E). Light-dependence profile of scotopic a- and b-wave amplitudes of Mut-*Rho^wt/hum^* mice treated with AAV-SaCas9/17-Sg2(blue) and AAV-SaCas9/CTRL (red) at 1×10^9^ dose (B, 3 mpi), 3×10^9^ dose (D, 3 mpi) and at 3×10^9^ dose (F, 11 mpi). Error bars show SEM, and the significance was calculated using two-tailed unpaired t-test, * *p*<0.05; ** *p*<0.01; *** *p*<0.005.

To evaluate whether silence or ablation of the *RHO*-5m allele could result in prevention or delay of photoreceptor cells loss, we evaluate the retinal structure in SaCas9/17-Sg2- and SaCas7/CTRL-treated Mut-*Rho^wt/hum^* mice at 4 mpi, 9 mpi and 11 mpi, at 1×10^9^ and/or 3×10^9^ dose (Supplementary Table S2). The average number of rows of ONL nuclei of SaCas9/17-Sg2-treated retinas was not significantly different from that of SaCas7/CTRL-treated retinas at 4 mpi (Supplementary Fig. S6A-S6D), either at 1×10^9^ or 3×10^9^ doses. Later at 9 mpi, either at 1×10^9^ or 3×10^9^ doses, measurement of the average number of rows of ONL nuclei revealed that there were significantly more photoreceptor cells in SaCas9/17-Sg2-treated retinas than in the age-matched untreated retinas (Fig. 9A-9C), IF staining showed that SaCas9/17-Sg2-treated Mut-*Rho^wt/hum^* mice had longer OS/IS (Fig. 9A-9B), which retained until at least 11 mpi (Fig. 9D-9E). These results showed clearly that SaCas9/17-Sg2 effectively restored the retinal function and preserved photoreceptor cells in the humanized RHO mtant mouse model, supporting the potential of SaCas9/17-Sg2 as an effective drug for RHO-T17M associated retinitis pigmentosa.

**Fig. 9.**
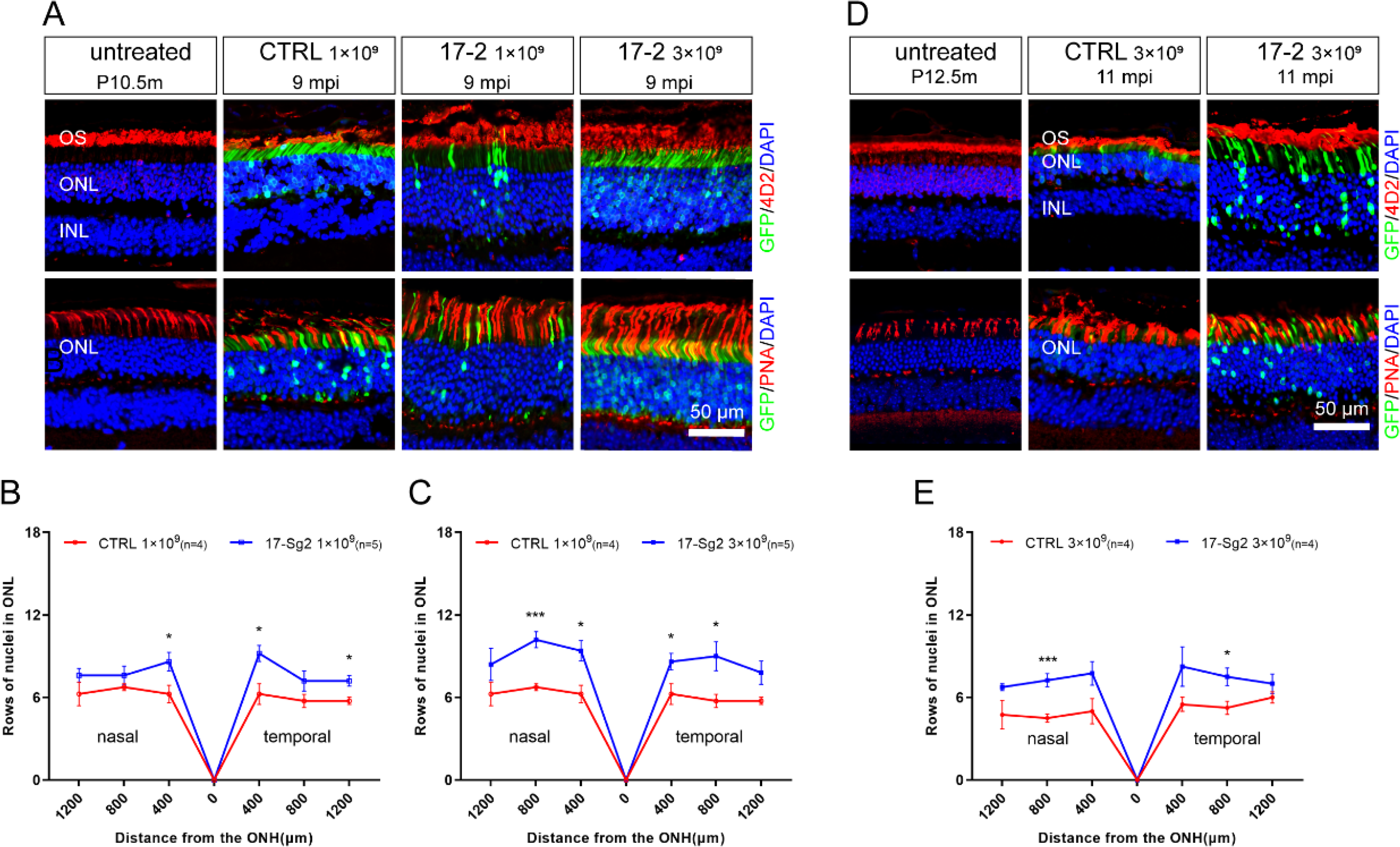
Photoreceptor cell preservation after gene editing with AAV-based SaCas9/17-Sg2 in Mut-*Rho^wt/hum^* mice. (A) 9 mpi, immunofluorescent (IF) staining of rhodopsin (4D2, red) and PNA (red) from age-matched untreated mouse retina, the AAV2/8-EFS-SaCas9-U6-CTRL (AAV2/8-SaCas9/CTRL)-treated and AAV2/8-SaCas9/17-Sg2-treated retinal regions (GFP+) in Mut-*Rho^wt/hum^* mice at 1×10^9^ and 3×10^9^. The graph revealed the average number of rows of ONL nuclei in AAV2/8-SaCas9/17-Sg2-treated and AAV2/8-SaCas9/CTRL-treated Mut-*Rho^wt/hum^* mice at 1×10^9^ (B) and 3×10^9^ (C) dose at 9 mpi. (D) 11 mpi, IF staining of rhodopsin (4D2, red) and PNA (red) from age-matched untreated mouse retina the AAV2/8-EFS-SaCas9-U6-CTRL (AAV2/8-SaCas9/CTRL)-treated and AAV2/8-SaCas9/17-Sg2-treated retinal regions (GFP+) in Mut-*Rho^wt/hum^* mice at 3×10^9^. (E) The graph revealed the average number of rows of ONL nuclei in AAV2/8-SaCas9/17-Sg2-treated and AAV2/8-SaCas9/CTRL-treated Mut-*Rho^wt/hum^* mice at 3×10^9^ dose at 11 mpi. Error bars show SEM, and the significance was calculated using two-tailed unpaired t-test, \**p*<0.05; ** *p*<0.01; *** *p*<0.005. Rhodopsin and PNA were indicated in red, nuclei were stained blue by DAPI. ONL = outer nuclear layer; OS = outer segment; IS =inner segment. Scale bar = 50 μm.

### Analysis of off-target activity

To evaluate the off-target activity of SaCas9/17-Sg2 in human gDNA, possible off-target sites obtained from the Benchling’s CRISPR Tool (Supplementary Table S4) were analyzed. The T7E1 assay showed no detectable off-target activity for these candidate sites for SaCas9/17-Sg2(OT1-OT10, Supplementary Fig. S7), which was consistent with the results of Sanger sequencing (data not shown). Besides, the top 20 off-target sites of 17-Sg2 were also predicted by the Cas-OFFinder online tool (Supplementary Table S5), comparing the next-generation sequencing (NGS) results of PCR amplicons amplified from 293T cells which were transfected with 17-Sg2 plasmid and CTRL plasmid, we observed no detectable off-target activity for these candidate sites (Supplementary Table S5). As illustrated in Supplementary Table S5, considering that SaCas9/17-Sg2 targeted *RHO*-T17M allele specifically, the corresponding *RHO*-WT sequence from 293T was supposed to a specifical off-target site, thereby the genomic region that was flanking the specifical off-target site was also amplified and sequenced, comparing the NGS results of 293T cells transfected with 17-Sg2 plasmid and CTRL plasmid, no detectable off-target activity for this site was detected. Together, these results confirmed the allele-specific gene editing activity of SaCas9/17-Sg2. To further examine the off-target potential in human gDNA resulting from SaCas9/17-Sg2, we performed whole-genome sequencing (WGS) in 293T cells transfected with 17-Sg2 plasmid and the control 293T cells (without transfection). We analyzed the single-nucleotide variants (SNVs) and indels in two cell groups. A total of 3,336,802 SNVs and 1,361,010 indels were detected in 17-Sg2 cells, which were not much different from the control (3,367,623 SNVs and 1,383,287 indels) (Fig. 10). We then filtered the SNVs and indels using Cas-OFFinder-predicted off-target sites and the WGS sequence for 17-Sg2 cells and untreated cells. We found 1 SNVs and 8 indels (Supplementary Table S6 and Table S7), but none of them were located in coding regions, suggesting no functional off-target sites in SaCas9/17-Sg2-treated human gDNA.

**Fig. 10.**
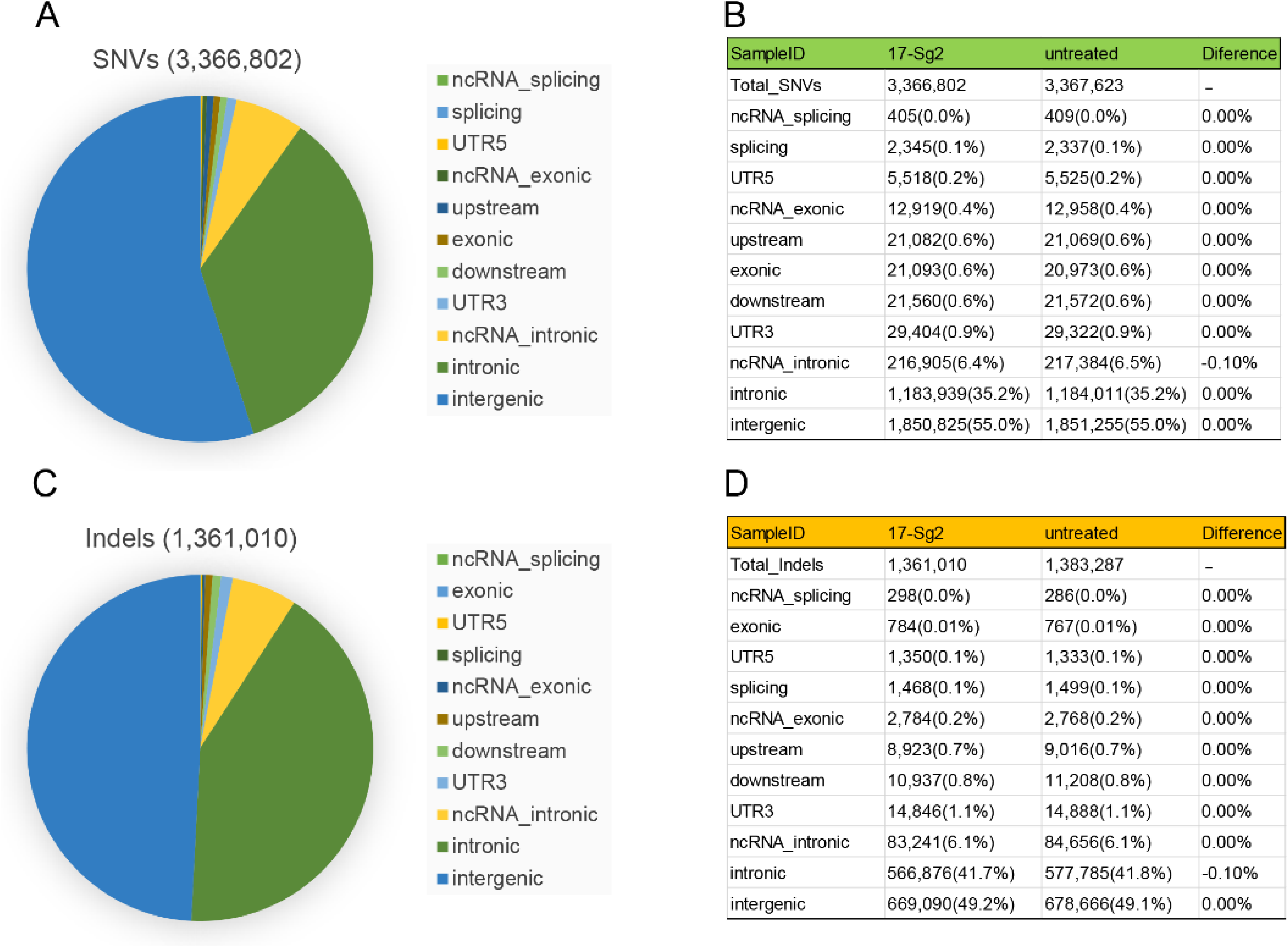
Examination of SaCas9/17-Sg2 off-target effects in human gDNA using whole-genome sequencing (WGS). Identification of single-nucleotide variants (SNVs, A) and indels (C) in 293T cells transfected with pX601-EFS-SaCas9-U6-17-Sg2 (17-Sg2) plasmid at the WGS level. The type of SNVs (B) and indels (D) in 293T cells transfected with 17-Sg2 plasmid and untreated cells at the WGS level.

## Discussion

There is currently an increasing number of gene editing clinical trials to treat hereditary diseases such as Sickle Cell Disease, β-Thalassemia(Frangoul et al., 2021) and transthyretin amyloidosis(Gillmore et al., 2021), and even cancer(Stadtmauer et al., 2020). These published data demonstrated the safety and feasibility of the CRISPR/Cas9 system *in vivo*. Previously, scientists have contributed significantly to the treatment of *RHO*-associated adRP using the allele-specific CRISPR/Cas9 gene editing strategy in iPSCs and animal models(Behnen et al., 2018, Bakondi et al., 2016), emonstrating the promising prospect of this approach in developing effective treatment for this disease. However, none of these methods can be directly applied in clinical practice. Herein, we developed an allele-specific gene editing therapeutic for *RHO* c.50C>T (p.T17M) for the first time. SaCas9/17-Sg2 was highly active and specific to the human *RHO* target sequence. *In vitro* experiments using 293T cells and patient-specific iPSCs have demonstrated strong nuclease specificity. A single AAV2/8 containing the SaCas9 and 17-Sg2 could be directly used as a therapeutic drug for adRP caused by the *RHO*-T17M mutation. This potential application was experimentally tested by subretinal delivery of AAV2/8-EFS-SaCas9-U6-17-Sg2 into the *RHO* humanized mice. Analysis of the treated retinas clearly showed its long-term therapeutic effect (up to 11 mpi) including retinal function improvement and photoreceptor preservation in the mutant humanized heterozygous mice(Liu et al., 2021a). Moreover, there was no unwanted off-target activities observed. Thus, our study makes a significant step towards the development of an effective drug for *RHO-*adRP.

Compared with allele-specific gene editing therapy, allele-independent gene ablation and replacement approaches are more cost-effective and efficient. However, there are potential risks with the gene ablation and replacement strategy. First, prior studies show that there is a fine balance between insufficiency and toxicity of rhodopsin (Lewin et al., 2014). Changes in the rhodopsin level (either suppression of rhodopsin expression or overexpression) may alter the morphology of ROS and discs(Makino et al., 2012, Wen et al., 2009). It is difficult for the ablation and replacement approach to strike such a balance *in vivo*. Second, previous studies have indicated that *Rho^-/-^* mice lose all of their rod photoreceptors extremely rapidly (approximately 3 months) (Humphries et al., 1997, Hobson et al., 2000), implying that disruption of both WT and mutant *RHO* alleles may aggravate the disease, when the exogenous augmentation of WT cannot rescue this disruption. Allele-specific silencing of mutant rhodopsin at the RNA or DNA levels can avoid the above problems. Although rhodopsin function is thought to be essential for the formation of rod OS and phototransduction initiation(Gunkel et al., 2015), heterozygous rhodopsin-knockout mice showed little retinal degeneration, indicating that one copy of this gene is sufficient for normal retinal development and functions(Lem et al., 1999). In this respect, maximal therapeutic benefit is expected when the mutant allele is completely ablated, while the normal allele is functionally intact. However, many factors may affect therapeutic effects. For instance, ASO can deregulate mutant rhodopsin levels transcriptionally, but the degree to which the toxic protein must be reduced to eliminate any disease-causing effect is not known(Shin et al., 2016). The presence of gain-of-function mutations and repeated subretinal injection may diminish the effectiveness of treatment. The CRISPR/Cas9 gene editing strategy in mammalian cells can provide a means to delete or correct disease-causing mutations while leaving the gene under the control of its endogenous regulatory elements(Jinek et al., 2013, Mali et al., 2013, Burnight et al., 2018), making this approach also promising for the treatment of inherited diseases.

CRISPR/Cas9 gene editing therapy is currently being advanced as the treatment of human IRDs(Maeder et al., 2019, Burnight et al., 2018). A study conducted by Maeder *et al*.(Maeder et al., 2019) shows that one of the complexities of drug development for gene editing is that it is human genome-specific. Herein, we used human cells and the *RHO* humanized mice to demonstrate that SaCas9/17-Sg2 could reach efficient and allele-specific gene editing, thereby improving the retinal functions and delaying photoreceptor cells loss. We also proved that SaCas9/17-Sg2 did not result in off-target activity using WGS. *In vitro* patient-derived iPSCs generated in this study offered as an ideal research material to validate gene editing specificity and cutting efficiency at the DNA level. Moreover, 3D retinal organoids or rod cell precursors derived from iPSCs (Capowski et al., 2019) have the potential to be used to mimic disease phenotypes or for autologous transplantation (Assawachananont et al., 2014, Aghaizu et al., 2017, Santos-Ferreira et al., 2016) after gene editing. However, we must emphasize that the transfection and gene editing efficiency in iPSCs are evidently lower than those of some commonly used cell lines,(Burnight et al., 2017, Lin et al., 2014) which may be due to the significant cell death that probably results from an overwhelming cytosolic dsDNA-induced innate immune response(Sun et al., 2013, Kim et al., 2018), implying that the on-target and off-target effects in iPSCs might be underestimated.

The first factor that may raise safety concerns for allele-specific CRISPR/Cas9 drug development is the specificity of SaCas9/SgRNA. This study demonstrated that SaCas9/17-Sg2 did not disrupt human WT *RHO* both *in vitro* and *in vivo*. In fact, a large number of allele-specific sgRNAs can be designed for disease-causing mutations in *RHO* (over 200 mutations, HGMD), such as the three sgRNAs for T17M. However, not all of them work well. A previous study proves theoretically and practically that the specificity of sgRNA depends on its seed sequence, which is a stretch of 8∼14 nt immediately upstream of the PAM. Even point mutations within the seed sequence can abolish cleavage by SpCas9 nucleases(Hsu et al., 2013, Jiang et al., 2013, Semenova et al., 2011), and spCas9 nuclease tolerates single-base mismatches in the PAM-distal region to a greater extent than in the PAM proximal region(Hsu et al., 2013, Jinek et al., 2012). Herein, the specificity of 17-Sg1 and Sg2 are better than that of 17-Sg3, may due to the fact that the point mismatch (c.50C>T) is within the seed sequence of 17-Sg1 and Sg2 but not within that of 17-Sg3, (Fig. 1), showing that the targeting specificity of SaCas9/SgRNA is also dependent on the seed sequence. Previous studies have indicated that T17M rhodopsin proteins fail to fold correctly, are retained in the ER and cannot easily reconstitute with 11-cis-retinal(Mendes et al., 2005, Behnen et al., 2018). Our experiments in Hela cells and 293T cells indicates rhodopsin protein with T17M and five variants are retained in cytoplasm and the endoplasmic reticulum(Liu et al., 2021a). Another factor may safety concerns is that indels created by gene editing possibly result in new rhodopsin variants which may also exert toxic effect or dominant-negative on WT rhodopsin(Patrizi et al., 2021). *In Silico* analysis and detailed *in vitro* experiments indicate that *RHO*-5m mRNA with these most frequent indels generated by gene editing near the N-terminal of *RHO* gene can be degraded due to the effect of NMD (Fig. 6 and Supplementary Fig. S5). These results support the potential safety of gene editing of the N-terminal domain of rhodopsin.

Currently, there are few available animal models compared to the number of known *RHO* mutations (over 200). Humanized animals are revolutionizing our ability to study and model disease and permit testing of the therapeutic effect of CRISPR/Cas9 in preclinical *in vivo* systems without putting humans at risk(Hosur et al., 2017). The shortage of humanized animal models may hamper the application of CRISPR/Cas9 technologies from animals towards patients. In the past, several *RHO* humanized mouse models were bred to express mutant human rhodopsin in a heterozygous knockout mouse rhodopsin (*Rho^+/-^*) or WT background (Lem et al., 1999, Chan et al., 2004, Olsson et al., 1992) by introducing human *RHO* genomic fragments that encompassed the entire transcriptional unit as well as its upstream and downstream flanking sequences. In contrast, our humanized models were generated by replacing the mouse *Rho* gene with the human *RHO* gene, but did not interrupt the flaking sequencing of this gene. Our study indicated(Liu et al., 2021a) that the typical adRP phenotypes of Mut-*Rho^wt/hum^* mice resulted from the *RHO*-5m allele, and the slow disease progression offered us the opportunity to observe therapeutic effects after treatment. Importantly, mice with different genotypes can be obtained from the intercross of these models according to our needs, thereby faciltating our *in vivo* experiments. The humanized mouse models combined with our strategy to selectively target the mutant *RHO* allele faithfully demonstrated their application prospects as well.

Recently, AAV has been considered an ideal choice for therapeutic gene delivery in the retina and has been confirmed to be promising for the delivery of CRISPR/Cas9 reagents(El Refaey et al., 2017, Yang et al., 2016). Besides Luxturna, many gene therapy clinical trials based on AAVs are ongoing (www.clinicaltrials.gov), indicating the feasibility and safety of AAVs. However, AAVs have a relatively small packaging capacity(∼4.7 kb), which limits the size and number of sequences that can be incorporated into a single virus(Burnight et al., 2018, Ran et al., 2015). To put all components into one AAV vector, the cytomegalovirus (CMV) enhancer promoter is replaced by the EFS promoter, and SaCas9, which is much smaller than SpCas9(Ran et al., 2015), is chosen and packed in an AAV vector with its sgRNA. As long as 9 mpi, no detectable toxic effects in mouse retinas were identified at three different doses when comparing the retinal structures of treated and untreated mice. Gene editing efficiencies herein are dose-dependent, yet the specificity of AAV-SaCas9/17-Sg2 is not affected by increasing AAV dose. However, indel analysis by Sanger sequencing in mouse retinas after treatment showed that a 49 bp insertion from the AAV inverted terminal repeat (ITR) sequence occurs at the cut site at a dose of 3×10^9^. Experiments in 293T cells showed that this insertion could lead to NMD of the *RHO*-5m mRNA. Previous studies confirmed that the majority of AAV genome insertions consist of an ITR sequence(Jarrett et al., 2017), which might be due to the combined properties of AAV ITR and its interaction with NHEJ host-cell machinery(McCullough et al., 2019, Young and Samulski, 2001). In this study, no AAV ITR insertions in treated retinas were detected in the 5×10^8^ and 1×10^9^ dose groups. We consider whether AAV dose will influence the possibility of insertions in question, and more research should be conducted to validate this hypothesis. At either DNA or RNA levels, the therapeutic effect in the retina at the 3×10^9^ dose was better than that at the 1×10^9^ dose. Previous studies(Mahfouz et al., 2011, Hsu et al., 2013) indicate that enzymatic specificity and activity strength are often highly dependent on reaction conditions, high enzyme concentration will amplify off-target activity, and one potential strategy for minimizing nonspecific cleavage is to limit the enzyme concentration. Therefore, the balance between therapeutic efficiency and AAV dose of subretinal injection *in vivo* must be weighted. Since approximately 10% of functional foveal cone photoreceptors are sufficient for near-normal visual acuity, researchers hypothesize that correction of the *CEP290* IVS26 mutation in at least 10% of foveal cones will lead to clinical benefit in patients(Maeder et al., 2019). The average cutting efficiencies in GFP+ retinas at 1×10^9^ or 3×10^9^ doses are more than 10%, and the long-term therapeutic effect in more treated mice should be evaluated to obtain more unbiased results.

In summary, this study demonstrates the safety and effectiveness of our allele-specific gene editing therapy *in vitro*and in humanized mouse models, thereby offering a generalizable framework for the preclinical development of gene editing medicine and future clinical trials.

## Material and Methods

### Animals

Approval was obtained from the Peking University Health Science Center Ethics Committee for Experimental Animal Research (Research License LA20200473). All procedures were performed according to the regulations of the Association for Research in Vision and Ophthalmology’s statement for the use of animals in ophthalmic and vision research. All mice were maintained in accordance with the guidelines of the Association for the Assessment and Accreditation of Laboratory Animal Care.

*RHO* humanized mouse models (with/without *RHO*-5m allele) with a C57BL/6J background using HDR and CRISPR/Cas9 strategy were custom designed and obtained from Beijing Biocytogen Co., Ltd (Beijing, China). Mice were bred and maintained at the Peking University Health Science Center Animal Care Services Facility under specific pathogen-free conditions with a 12 h light/12 h dark cycle. Food and water were provided *ad libitum*.

### Patients

The study was approved by the Medical Scientific Research Ethics Committee of Peking University Third Hospital (Research License 2021262). The procedures were performed in accordance with the tenets set forth in the Declaration of Helsinki. All patients provided written informed consent for this study.

### Generation of patient-specific iPSCs

Urine samples from a male patient (45-year-old) affected with adRP resulting from *RHO-*T17M mutation(Liu et al., 2021b) and a healthy male control (27-year-old) were collected. Urine-derived cells (UCs) were isolated using a UC isolation medium (CA3102500, Cellapy) and cultured for two passages in a UC expansion medium (CA3103200, Cellapy). UCs were reprogrammed using the integration-free CytoTune™ 2.0 Sendai Reprogramming Kit (A16517, Thermo Fisher Scientific, MA, USA), which contained four Yamanaka factors, SOX2, OCT3/4, KLF4, and c-MYC, according to the manufacturer’s instructions. After the emergence of colonies with iPSC-like morphology, R1 medium (85851, STEMCELL) was used for the culture at 37 °C under 5% CO2 conditions. The primary antibodies and corresponding secondary antibodies for IF staining were listed below(Zhou et al., 2012, Takahashi et al., 2007, International Stem Cell et al., 2007).

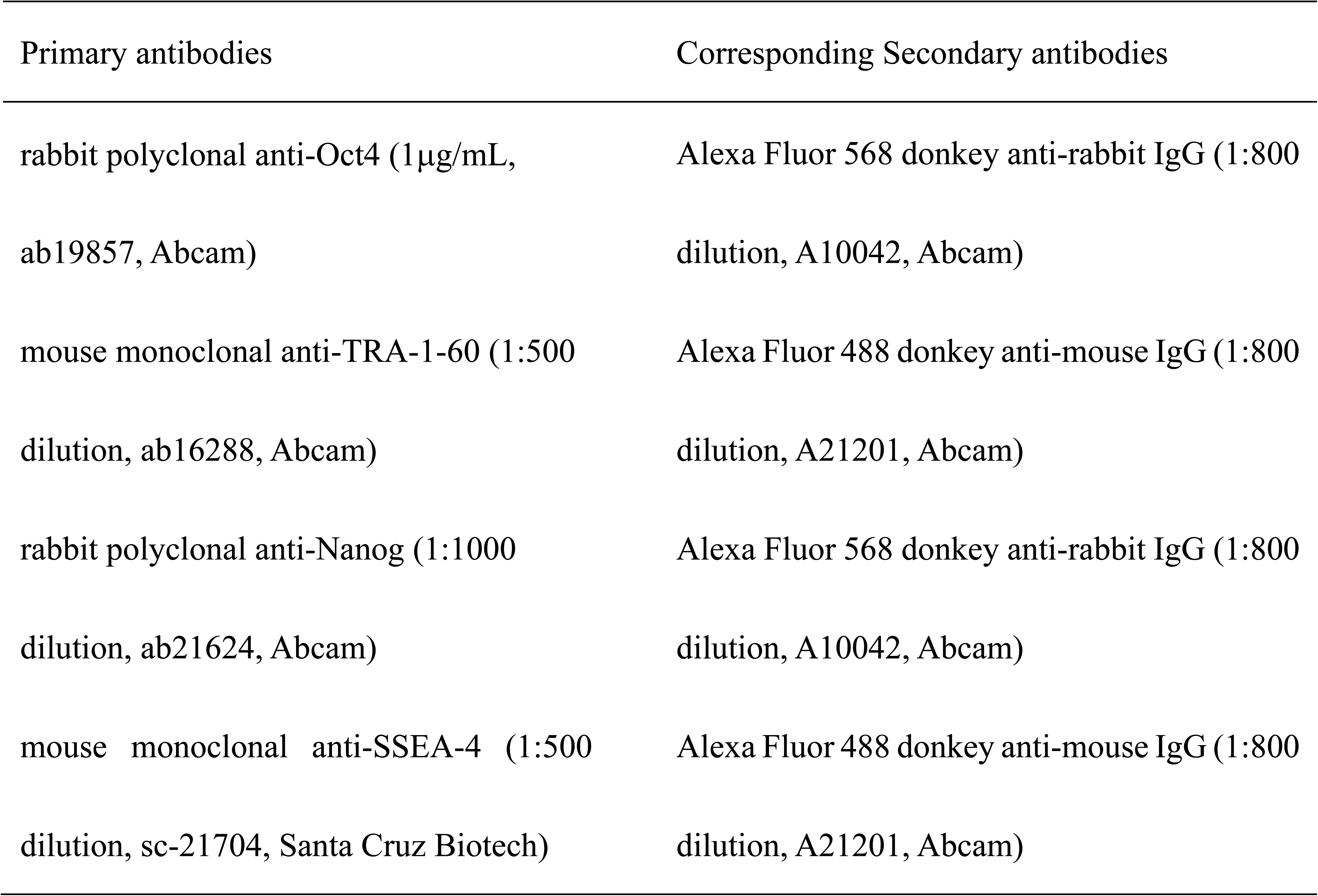

### SgRNA design and *in v*itro cleavage efficiency study

Oligo pairs of 21-nucleotide (nt) sgRNA sequences targeting the *RHO-*T17M mutation were designed based on the website https://www.benchling.com/. Based on the *RHO* gene sequence (ENSG00000198947), we identified all the NNGRR PAMs for SaCas9. Oligonucleotides corresponding to the sgRNA of interest were purchased from Beijing Ruibio Biotech Co., Ltd. (Beijing, China).

An *in vitro* cutting assay using the SaCas9/SgRNA Cutting Efficiency Detection Kit (VK012, Viewsolid Biotech, Beijing, China) was performed to ensure the relative cutting efficiency and specificity of sgRNAs. The sgRNAs of interest were directly transcribed *in vitro* with T7 polymerase from double-stranded DNA templates. Primer pairs RHO17-F and R (Supplementary Table S8) were used to amplify the targeted region with or without c.50C>T mutation, yielding a 678 bp PCR fragment, respectively. Afterwards, 1 μL of purified PCR amplicon (50 ng/μL), 1 μL of *in-vitro*-transcribed sgRNA (50 ng/μL), 1 μL of SaCas9 protein (1 U/μL), 2 μL of 10×SaCas9 buffer, and 15 μL of ddH2O were added into a tube and incubated for 30 min at 37 ℃. Subsequently, 2 μL of DNA loading buffer (9157, TAKARA, Kusatsu, Japan) was added to the reaction and incubated at 65 ℃ for 5 min. The full reaction was run on 2% agarose gel (111860, Biowest).

### Vector Constructs

The selected sgRNA was cloned into the plasmid pX601-AAVCMV: NLS-SaCas9-NLS-3xHA-bGHpA; U6: BsaI-sgRNA (pX601, #61591, Addgene). The CMV enhancer promoter was replaced by a commonly used short, ubiquitous promoter-shortened elongation factor alpha (EFS). The plasmid pX601-EFS-SaCas9 was linearized using BsaI restriction enzyme (R3733S, New England Biolabs, MA, USA), followed by purification with a gel extraction kit (D2500, Omega). The sgRNA of interest was cloned into the BsaI site using T4 DNA ligase (M0202S, NEB), and then transformed into Top10-competent bacteria. Clones were amplified in a liquid Luria-Bertani (LB) broth, followed by the extraction of plasmid DNA (D6943; OMEGA). In addition, the plasmid containing the puromycin selection cassette AAV-EFS-SaCas9-p2a-Puro was generated from pX601-EFS-SaCas9, and the selected sgRNA was cloned into this plasmid or into Lenti_SaCRISPR_GFP (ATCC, #118636). In replacing the SaCas9 sequence with the EGFP sequence, the pX601-EFS-GFP plasmid was generated.

For the rhodopsin overexpression experiments, a full-length *RHO* cDNA with or without the T17M mutation was cloned into a lentiviral expression vector FUGW (Addgene, # 14883), and the *RHO* CDS was inserted under the control of the UbC promoter. The full-length *RHO* cDNA with or without the T17M mutation was also cloned into the pmCherryN1 and pEGFPN1 vectors using the pEASY^®^-Basic Seamless Cloning and Assembly Kit (CU-201, TransGen Biotech, Beijing, China) according to the manufacturer’s protocol. *RHO* variant plasmids (*RHO*-c.46delG/c.45_46delTG/c.43_46del4bp/c.46_47ins49bp) based on the *RHO*-T17M-pEGFPN1 or *RHO*-5m-pEGFPN1 were generated. The primer pairs used were listed in Supplementary Table S8.

### Viral production

For the production of lentivirus, 293T cells were transfected with a combination of three plasmids, FUGW-*RHO*-cDNA or Lenti_SaCRISPR_GFP, Pax2, and vesicular stomatitis virus G protein (VSV-G) plasmid using polyetherimide (PEI) (B600070, ProteinTech Group, Chicago, USA) according to the manufacturer’s protocol. Twenty-four hours after transfection, the medium was replaced with a fresh one. Forty-eight hours later, the medium containing the viral particles was collected.

The AAV2/8 vectors that encoded the SaCas9 gene under the control of the EFS promoter and the sgRNA of interest under the control of the U6 promoter (AAV2/8-EFS-SaCas9-U6-SgRNA) were produced in 293T cells by PEI co-transfection with three different plasmids: pX601-EFS-SaCas9-U6-sgRNA plasmid (SgRNA plasmid), packaging plasmid pAAV-RC8 for the *rep* and *cap* genes, and the pHelper plasmid (Cell Boil, Inc). Furthermore, the AAV2/8-EFS-EGFP from the pX601-EFS-EGFP plasmid and the AAV2/8-EFS-SaCas9-U6-CTRL plasmid from the pX601-EFS-SaCas9-EFS-CTRL plasmid were also produced. The titers were determined by qPCR.

### Cell cultures and plasmids transfection or viral infection

HEK293T cells (CRL-3216, ATCC) were cultured in Dulbecco’s modified Eagle’s medium (DMEM, CM15019, Macgene, Beijing, China) and 10% fetal bovine serum (FBS, 10099-141C, Gibco, USA). For viral transduction, the medium was removed and replaced with a fresh one as well as the previously purified lentivirus. On the day after transduction, the cells were washed with 1×DPBS (CC010, Macgene, Beijing, China) and cultured in fresh medium. At the 72 h of transduction, infected 293T cells were transfected with plasmids using PEI. The fluorescence intensities of GFP and mCherry were examined using the Spark 10M Multiscan Spectrum (TECAN, Switzerland).

293T cells were transfected with WT, T17M, *RHO*-5m-pEGFPN1, or *RHO*-variants-pEGFPN1 plasmids using PEI. After 24 h or 72 h of transfection, GFP was observed using a Nikon fluorescence microscope (Nikon, Japan). After one week of transfection, the cells were fixed in 100% methanol (chilled at –20 °C) at room temperature for 5 min, incubating the samples were incubated for 10 min with PBS containing 0.1% Triton X-100. Cells were incubated with 5% normal donkey serum (017-000-121, Jackson Laboratory, USA) in PBST for 30 min at room temperature (RT) to block non-specific binding of the antibodies. The primary antibody used was monoclonal mouse monoclonal anti-rhodopsin (4D2) antibody (1:1000 dilution, ab98887, Abcam, Cambridge, MA)). The secondary antibody used was donkey anti-mouse antibody conjugated with Alexa Fluor 568 (1:800 dilution, A10037, Invitrogen). Transfected cells were counterstained with DAPI (1:5000 dilution, C0060, Solarbio, Beijing, China) and examined under a Nikon *n-Storm* confocal microscope (Nikon, Japan).

On the day of transfection, fresh R1 medium containing 10 μM Y-27632 (ROCK inhibitor, STEMCELL, Canada) was added to the iPSCs. iPSCs from the healthy male donor and the male adRP patient were transfected with AAV-EFS-SaCas9-U6-sgRNA-p2a-Puro plasmid using Lipofectamine^TM^ Stem Reagent (STEM00003, Thermo Fisher Scientific) according to the manufacturer’s protocol. For drug selection, R1 medium containing 0.1∼0.2 μg/mL puromycin (A1113803, Life, USA∼15 d) was added after 24 h of transfection.

### Subretinal injection

*RHO* humanized mice were treated on the postnatal month 1.5∼1.8 (P1.5∼1.8 m) with AAV vectors of AAV2/8-EFS-GFP or a 1:1 mixture of AAV2/8-EFS-GFP and AAV2/8-EFS-SaCas9-U6-sgRNA or a 1:1 mixture of AAV2/8-EFS-GFP and AAV2/8-EFS-SaCas9-U6-CTRL at different doses via subretinal injection in both eyes as described previously(Hu et al., 2020, Wu et al., 2015). A summary of mouse experiments was provided in Supplementary Table S2. Eyes with severe hemorrhage or leakage of the vector solution from the subretinal space into the vitreous were excluded from further study.

### Cell and tissue collection

At 3 mpi, two retinas from each mouse were dissected under a Nikon fluorescence microscope (Nikon, Japan). GFP+ and GFP-areas of the treated retinas were cut for further analysis, including genomic DNA (gDNA), RNA, and protein isolation. Genomic DNA was extracted from mouse retinas or cells and isolated using a DNA isolation kit (DC102, Vazyme, Nanjing, China). RNA was extracted and isolated from mouse retinas or cells using RNA isolation (RC101, Vazyme, Nanjing, China). In addition, gDNA and RNA were also extracted using TRIzol (15596026, Invitrogen, USA).

### On-target and off-target analysis

The genomic regions flanking the sgRNA target sites were amplified by PCR using 50-100 ng of gDNA as a template. Primers used were listed in Supplementary Table S8. For T7EI analysis, 500 ng of purified PCR products (D2500, Gel Extraction Kit, OMEGA, USA) were denatured and reannealed in NEB buffer 2 (M0302S, NEB, USA): 95 ℃, 2 min; 95 to 85 ℃ at –2 ℃/s; 85 to 25 ℃ at –0.1 ℃/s. Upon reannealing, 10 U of T7EI ((M0302S, NEB, USA) was added and incubated at 37 ℃ for 30 min. Afterwards, was analyzed on 2.5% agarose gels using a Gel Doc imaging system (Bio-Rad, USA). The amplicons obtained previously were also subjected to clone sequencing for identification of on-target indels. This consisted of cloning the PCR products using the ZTOPO BLUNT/TA with the Zero Background cloning kit following the manufacturer’s instructions (ZC211-2, ZOMANBIO, Beijing, China). In addition, PCR products for targeted deep sequencing were prepared as described in the Hi-TOM Kit19 (Novogene, Beijing, China). Meanwhile, mixed samples were sequenced on the Illumina HiSeq platform (www.hi-tom. net/hi-tom/documentation.html)(Hu et al., 2020, Liu et al., 2019).

For off-target analysis, 293T cells were transfected with sgRNA plasmid or CTRL plasmid using PEI. One week after transfection, gDNA was extracted. Off-target sites were obtained from the Benchling’s CRISPR Tool. The genomic regions that were flanking off-target sites were amplified by PCR and then validated using the T7E1 assay and Sanger sequencing. In addition, off-target sites were identified using the Cas-OFFinder online tool (http://www.rgenome.net/cas-offinder) (Bae et al., 2014). PCR amplicons were analyzed by NGS using the Illumina NovaSeq platform (Illumina, San Diego, California, USA). Furthermore, gDNA of 293T cells with or without transfection was tested by WGS using the Illumina NovaSeq platform (Illumina, San Diego, CA, USA).

### RNA analysis

Five hundred nanograms of RNA was used for cDNA synthesis using the One-Step gDNA Removal and cDNA Synthesis Supermix Kit (AT311, TransGen Biotech) at a final volume of 20 μL and cDNA was then diluted by adding 80 μL of RNase-free H2O. qPCR reactions were performed using the ABI7500 Real-Time PCR Detection System (Carlsbad, CA, USA). Each 20 μL reaction contains 10 μL of TransStart Top Green qPCR SuperMix (AQ132, TransGen Biotech), 0.2 μM of each forward and reverse primer, and 1 μL diluted cDNA. All reactions were performed in triplicates. The oligonucleotide sequences are listed in Supplementary Table S8.

### WB analysis

Cells or mouse retinas were homogenized in 200 μL of RIPA buffer (C1053+, Applygen, Beijing, China plus protease inhibitors). Protein quantification was performed using a bicinchoninic acid (BCA) kit (P1513, Applygen, Beijing, China). 10μg of Protein extract loaded onto a 10% sodium dodecyl sulfate-polyacrylamide gel. Following electrophoresis, proteins were transferred onto a PVDF membrane. The membrane was blocked with 5% milk in TBST (B1009, Applygen, Beijing, China) and incubated with a primary mouse monoclonal 1D4(1:1000 dilution, ab5417, Abcam, Cambridge, MA)), rabbit Anti-CRISPR-Cas9 antibody (1:5000 dilution, ab203943, Abcam, Cambridge, MA)), mouse monoclonal anti-β-actin antibody (1:5000 dilution, AF0003, Beyotime, Shanghai, China), or mouse monoclonal anti-GAPDH antibody (1:5000 dilution, ZL9002, ProteinTech Group, Chicago, USA) in primary antibody diluent for western (C1240, Applygen, Beijing, China) overnight at 4 °C, respectively. The next day, after washing 3 times with 1×TBST (B1109, Applygen, Beijing, China), the membrane was incubated with horse radish peroxidase (HRP)-conjugated Goat anti-mouse IgG antibody (A0216, Beyotime, Shanghai, China) or Goat anti-rabbit IgG antibody (A0208, Beyotime, Shanghai, China) at 1:1000 dilution for 1 h at RT. GAPDH or β-Actin was used as a loading control. Blots were analyzed using a ChemiDoc™ MP imaging system (Tanon Science & Technology Co., Shanghai, China).

### Histological analysis and morphological measurements

For cryosectioning, eyes were enucleated and fixed with 4% paraformaldehyde in PBS or FAS eyeball fixative solution (G1109, Servicebio, Beijing, China), the lens and cornea were removed, and the eye cups were embedded in optical cutting temperature compound (OCT, 4583, Tissue-Tek, Sakura Finetek, Torrance, CA) and frozen in liquid nitrogen. The eye cup was sectioned to a thickness of 7 μm using a cryostat. For histological analysis, sections were collected at regular intervals from approximately 24 sites per eye. Sections were stored at –80 °C and used within 2 to 3 days.

Before use, the cryosections were fixed on slides for 10 min with acetone. The sections were then washed thrice with PBS, followed by IF staining. The primary antibodies used were monoclonal mouse 4D2 antibody and biotinylated peanut agglutinin (PNA; 1:200 dilution, B-1075, Vector, CA, USA). The secondary antibodies used were donkey anti-mouse conjugated with Alexa Fluor 568 for 4D2, and rhodamine (TRITC)-conjugated streptavidin (1:200 dilution, 123126, Jackson ImmunoReseach Laboratories, West Grove, PA, USA) for PNA. 4D2 and PNA were used to identify the rod and cone photoreceptors, respectively. Cryosections were also stained with hematoxylin and eosin (HE), and whole slide images were obtained using a NanoZoomer digital slide scanner (Hamamatsu Photonics, Hamamatsu City, Japan).

### Full-field ERG

The ERG responses of mice (*n*≥3) were recorded using the Espion E^2^ recording system (Diagnosis LLC, Lowell, MA, USA). As shown in Supplementary Table S2 and our previous study (Liu et al., 2021a), the ERG recordings were initially conducted at 3 mpi under dark-(scotopic) and light-adapted (photopic) conditions.

### Quantification and statistical analysis

The results were evaluated using two-tailed unpaired Student’s t-test and are presented as the mean ± SEM. GraphPad Prism 9 (GraphPad Software, La Jolla, CA, USA) was used for statistical analyses. Statistical significance was set at *p* < 0.05.

## Acknowledgement.

The author thanks Dr. Juan Du for the gift of the AAV-EFS-SaCas9-p2a-Puro plasmid. This article was supported by the National Natural Science Foundation of China (grant nos.: 81770966) and the Beijing Natural Science Foundation of China (grant nos.: 19JCZDJC64000(Z)); The sponsors and/or funding organization had no role in the design or conduct of this research.

## Author contributions

L.X., and Y.L. planned the research; L.X., J.R. Q.J. and Z.F. performed the experiments; L.X. and J.R. analyzed the data; H.S. and M.X. contributed technical support; L.X., and Y.L. wrote the paper.

## Data Availability

The data that support the findings of this study are available from the corresponding author on reasonable request.

## Supplementary files

**Supplementary Fig. S1.**
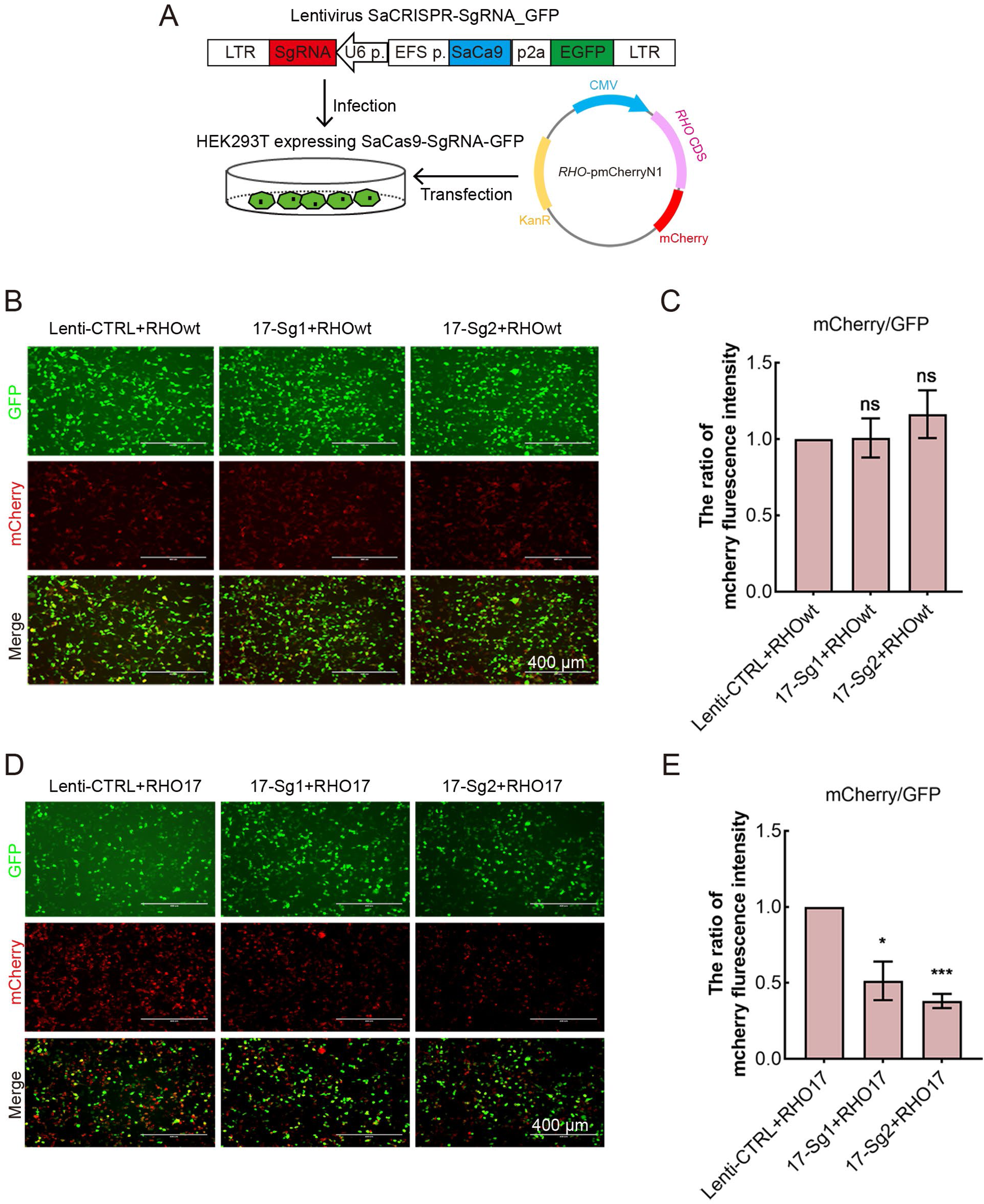
The knock-down of human *RHO*-T17M gene expression in 293T cells stably expressing human *RHO*. (A) SaCas9/SgRNA _GFP stable cell line generation of 293Tand transfection of human *RHO* cDNA plasmid. The calculation of ratio of mCherry/GFP fluorescence intensity (mCherry/GFP ratio) indicated SaCas9/SgRNA did not cut the wildtype RHO allele(B-C). While SaCas9 with 17-Sg1(SaCas9/17-Sg1) and SaCas9/17-Sg2 could significantly reduce mCherry/GFP ratio (D-E). Scale bar=400μm. Error bars show SEM, and the significance was calculated using two-tailed paired t-test, ns indicated no significance, \**p*<0.05, *** *p*<0.005.

**Supplementary Fig. S2.**
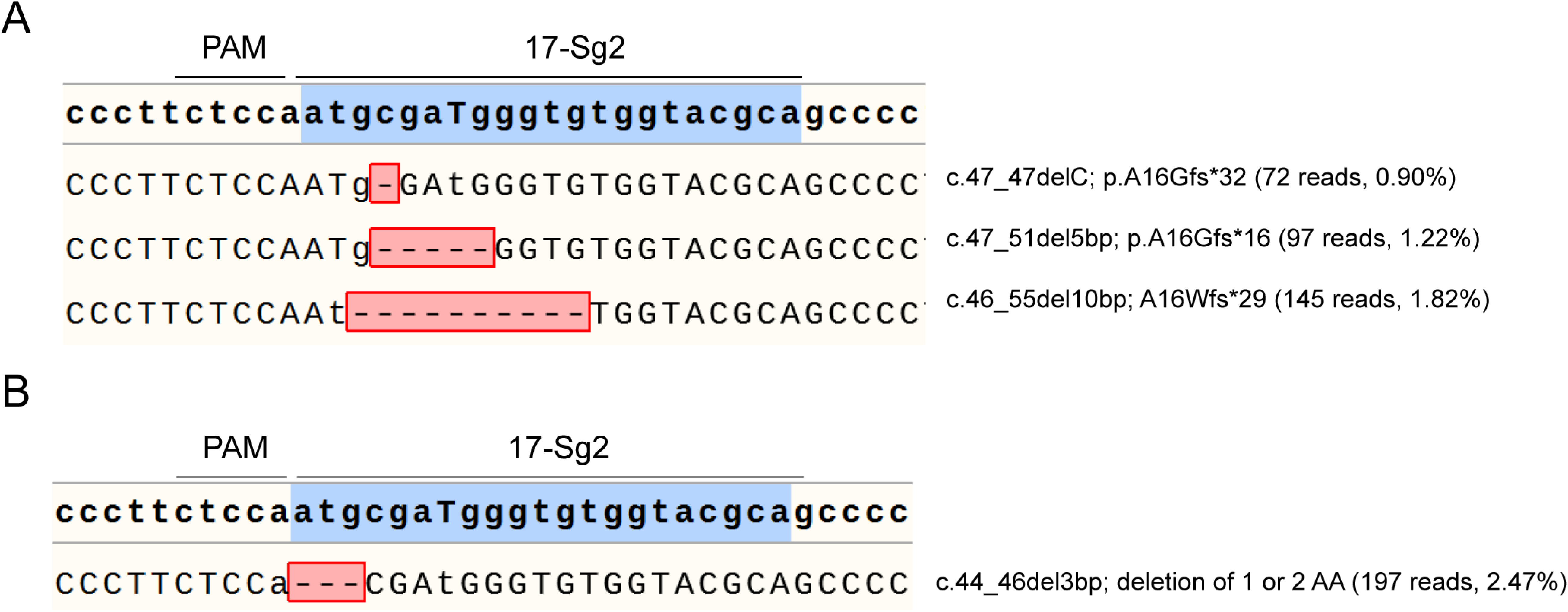
Hi-Tom sequence results of SaCas9 and 17-Sg2-treated P1 iPSCs(A) and P1 iPSCs colony(B).

**Supplementary Fig. S3.**
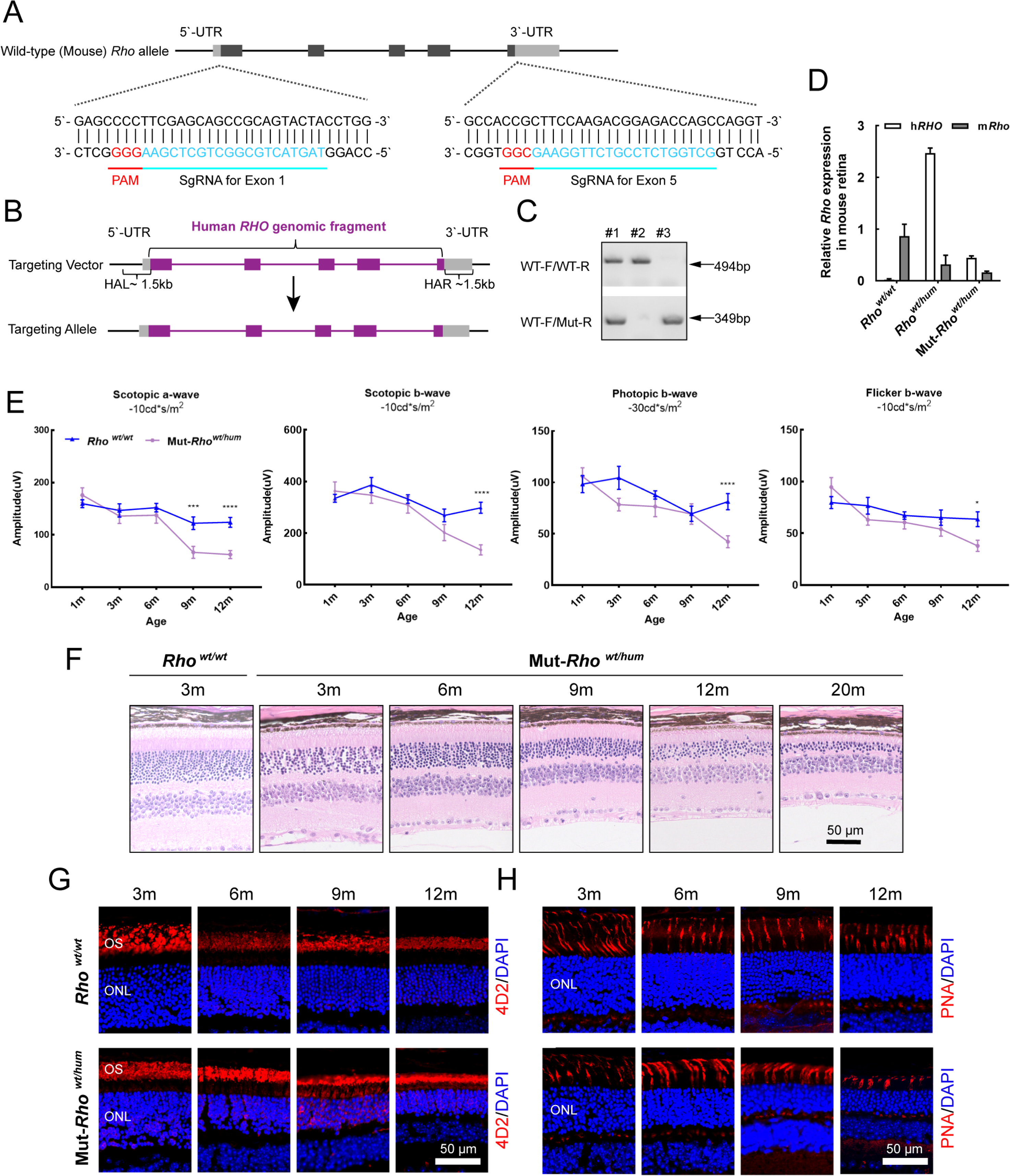
Generation and phenotypes of the humanized mouse models. (A) Schematic representation of the two Cas9/gRNA-targeting sites in exon one and exon five of mouse *Rho* gene, respectively. Targeted genomic sites are indicated in blue. Protospacer adjacent motif sequences were marked in red. Exons were indicated by closed boxes. (B) Schematic representation of the targeting vector containing the replacement of mouse exon1(mE1) after the start codon, mE2, mE3, mE4, mE5 before the stop codon, as well as mouse intron1(mI1), mI2, mI3 and mI4, by the human counterparts and the humanized targeting allele. (C) Scotopic and photopic ERG results of *Rho ^wt/wt^* and Mut-*Rho ^wt/hum^* mice with increasing age. Error bars show SEM, and the significance was calculated using two-tailed paired t-test, \**p*<0.05; ** *p*<0.01; *** *p*<0.005. (D) Histologic sections of Mut-*Rho ^wt/hum^* mice and WT controls. A gradual loss in the number of photoreceptor cells in the ONL was identified with age in Mut-*Rho ^wt/hum^* mice. Scale bar = 50μm. Immunofluorescent staining of rhodopsin (E), PNA (F) in Mut-*Rho ^wt/hum^* mice and WT controls. Rhodopsin (4D2 antibody) and PNA staining were shown in red. Nuclei were stained blue by DAPI. Scale bar = 50μm, OS = outer segment, ONL = outer nuclear layer.

**Supplementary Fig. S4.**
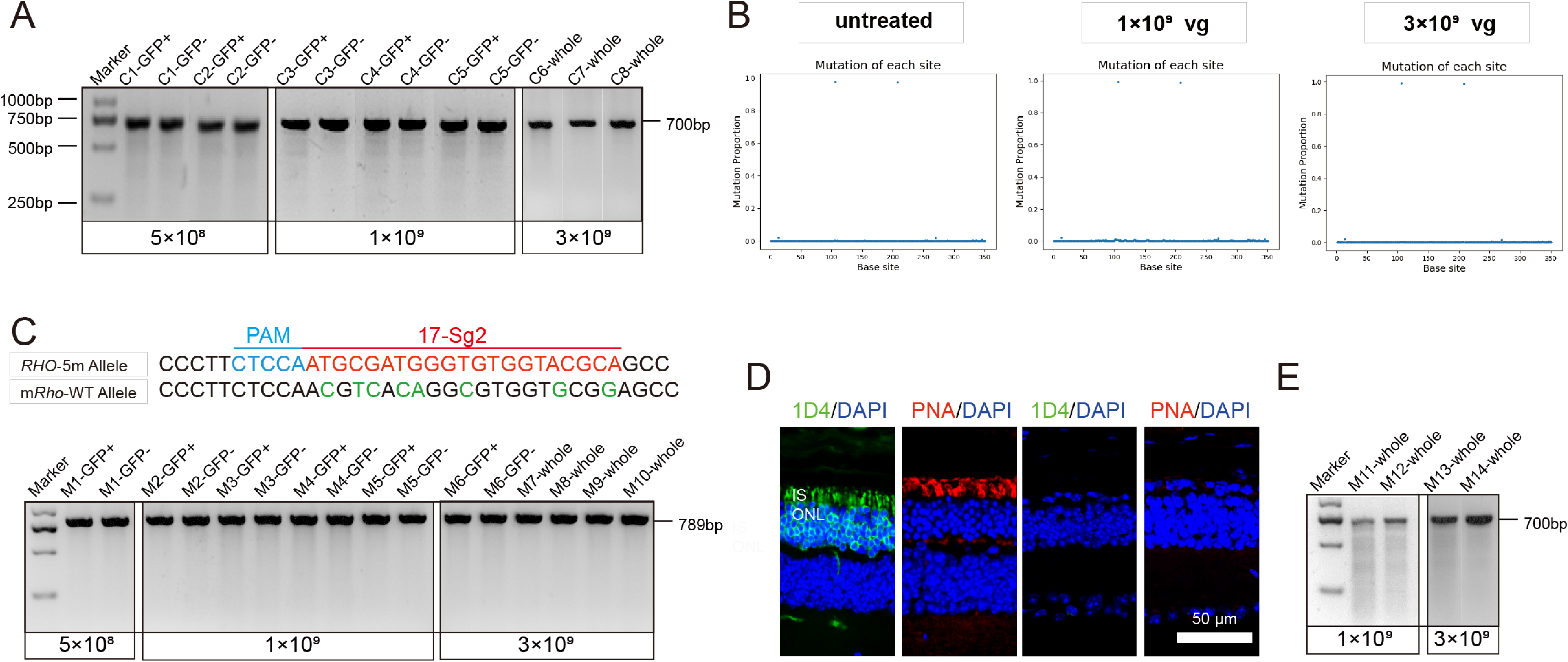
Analysis of gene editing specificity and efficiency in *RHO* humanized mouse retinas. (A) T7E1 assay performed on GFP+ and GFP-retinas indicated undetectable gene editing activity (gene editing specificity) of AAV2/8-EFS-SaCas9-U6-17-Sg2 (AAV2/8-SaCas9/17-Sg2) in *Rho^wt/hum^* retinas at 5×10^8^, 1×10^9^ and 3×10^9^ doses, the full-length amplicon was 700bp. (B) PCR-based Next-generation sequencing indicated AAV2/8-SaCas9/17-Sg2 did not cut wild type human *RHO* (h*RHO*-WT) allele in *Rho^wt/hum^* retinas at 1×10^9^ and 3×10^9^ doses and untreated *Rho^wt/hum^* retinas, the X axis represented the base site, the Y axis represented the indels proportion in h*RHO*-WT allele of AAV2/8-SaCas9/17-Sg2-treated *Rho^wt/hum^* retinas. (C) T7E1 assay performed on GFP+ and GFP-retinas indicated that AAV2/8-SaCas9/17-Sg2) in did not cut WT mouse *Rho* in allele Mut-*Rho^wt/hum^* retinas at 5×10^8^, 1×10^9^ and 3×10^9^ doses, the full-length amplicon was 789bp, 17-Sg2 was marked in red; Protospacer adjacent motif sequence was marked in light blue and mismatches in mouse Rho sequence comparing to 17-Sg2 were indicated in green. (D) IF staining indicated rapid loss of photoreceptor cells in Mut-*Rho ^hum/hum^* mice, rhodopsin staining (1D4) was shown in green, PNA staining was shown in red, nuclei were stained blue by DAPI. Scale bar = 50 μm. IS = inner segment, ONL=outer nuclear layer. (E)T7E1 assay indicated undetectable cutting activity in AAV2/8-SaCas9/17-Sg2-treated Mut-*Rho^hum/hum^* retinas at 1×10^9^ dose and 3×10^9^ dose.

**Supplementary Fig. S5.**
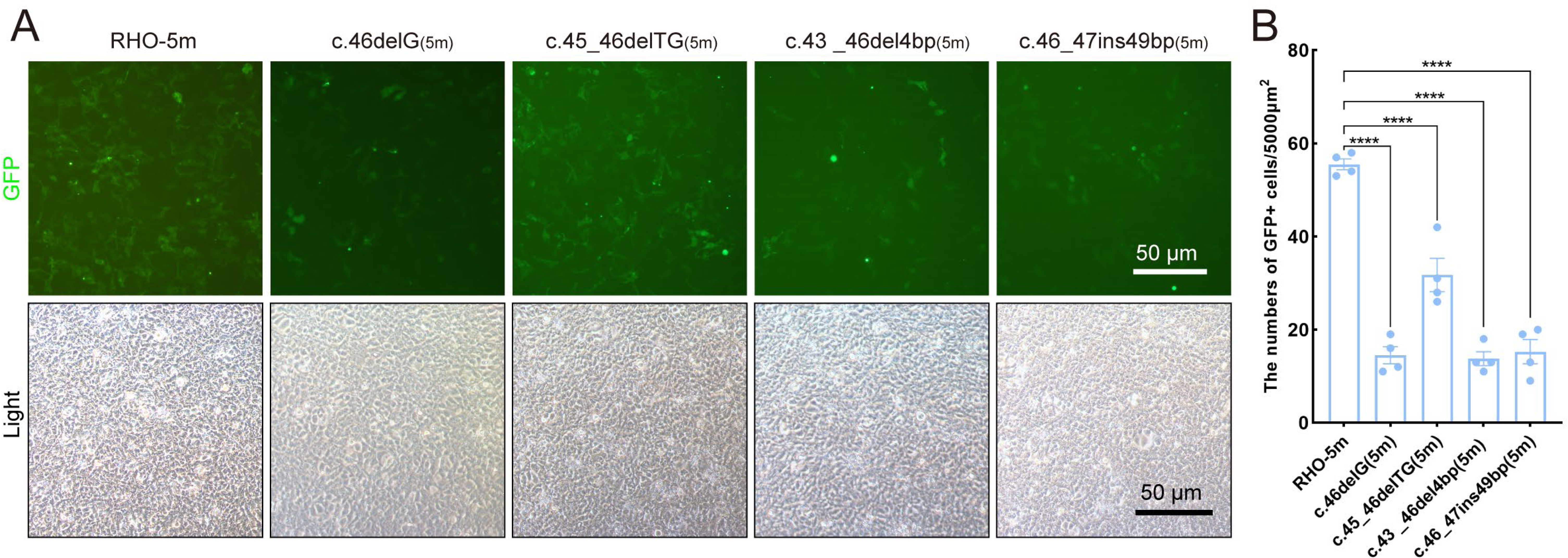
Expression of the mutant human *RHO* allele after gene editing with SaCas9/17Sg-2 *in vitro*. (A-B) GFP expression in 293T cells transfected with pEGFPN1 vector carrying *RHO*-5m (five variants in *RHO* CDS) and four edited variants, 24 hours after transfection, Scale bar = 50μm.

**Supplementary Fig. S6.**
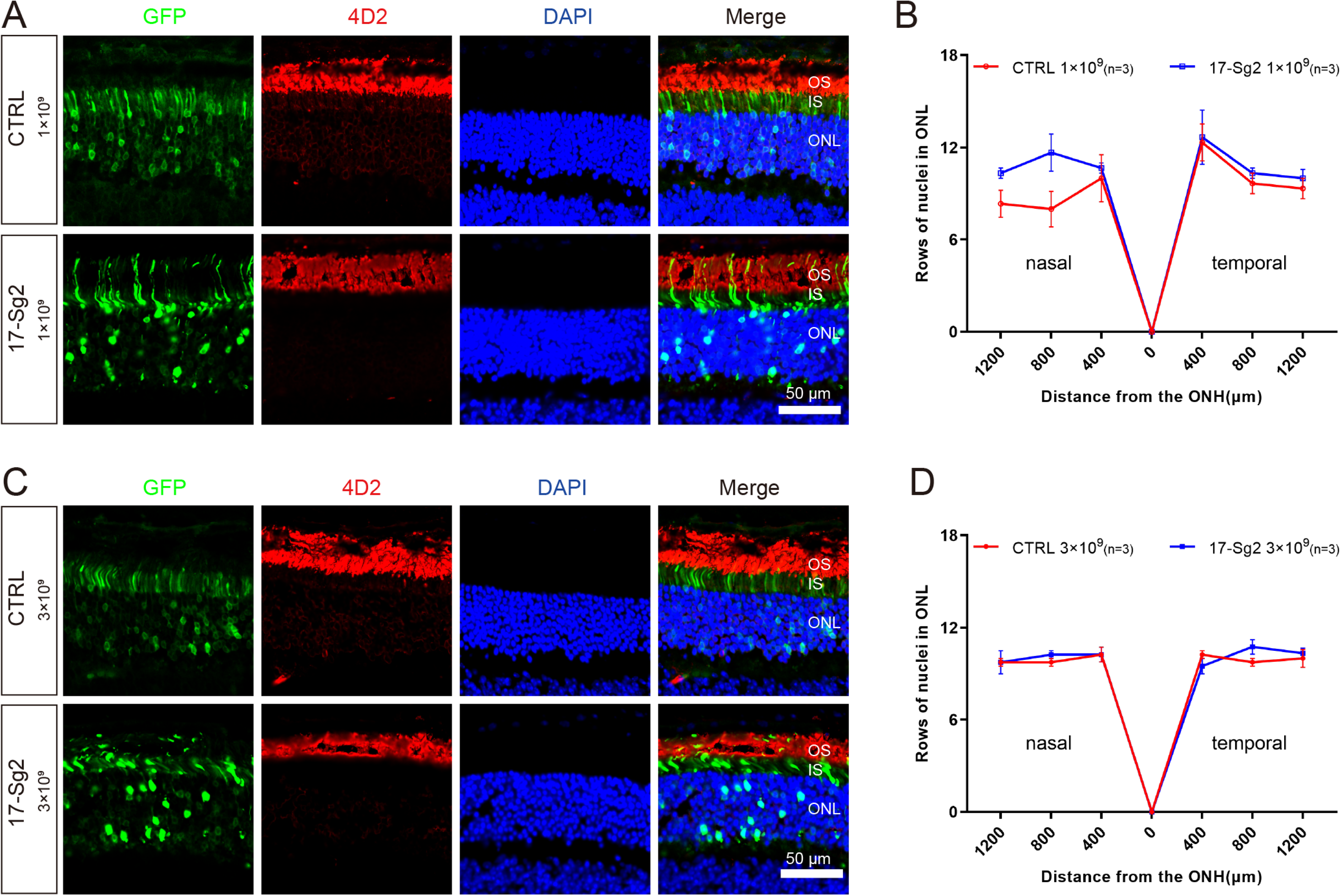
Retinal images after gene editing with AAV-based SaCas9/17-Sg2 in Mut-*Rho^wt/hum^* mice. Retinal images from the treated (GFP+) regions in Mut-*Rho^wt/hum^* mice at 1×10^9^ dose (A) and 3×10^9^ dose (C). (Upper) Representative image of mouse retina treated with AAV2/8-EFS-SaCas9-U6-CTRL (AAV2/8-SaCas9/CTRL); (Lower) Representative image of mouse retina treated with AAV2/8-SaCas9/17-Sg2. The graph reveals the average number of rows of ONL nuclei in SaCas9/17-Sg2-treated mice and SaCas9/CTRL-treated mice at 1×10^9^ dose (B) and 3×10^9^ dose (D) at 4 mpi. Error bars show SEM, and the significance was calculated using two-tailed unpaired t-test. Rhodopsin (4D2) was in indicated in red, nuclei were stained blue by DAPI. ONL= outer nuclear layer; OS = outer segment; IS= inner segment. Scale bar = 50 μm.

**Supplementary Fig. S7.**
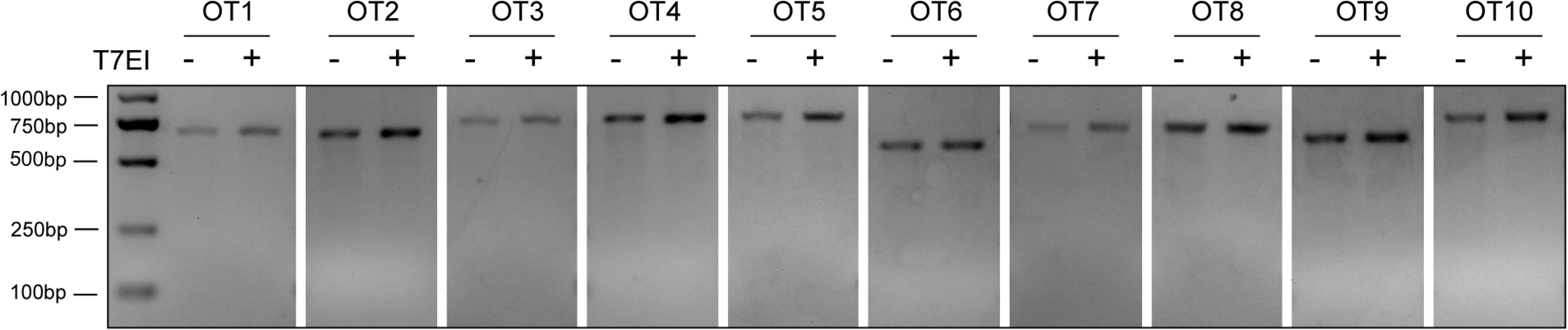
Analysis of SaCas9/17-Sg2 off-target activity. T7E1 assay showed undetectable off-target activity for all candidate sites for 17-Sg2 (OT1-OT10).

**Supplementary Table S1.**
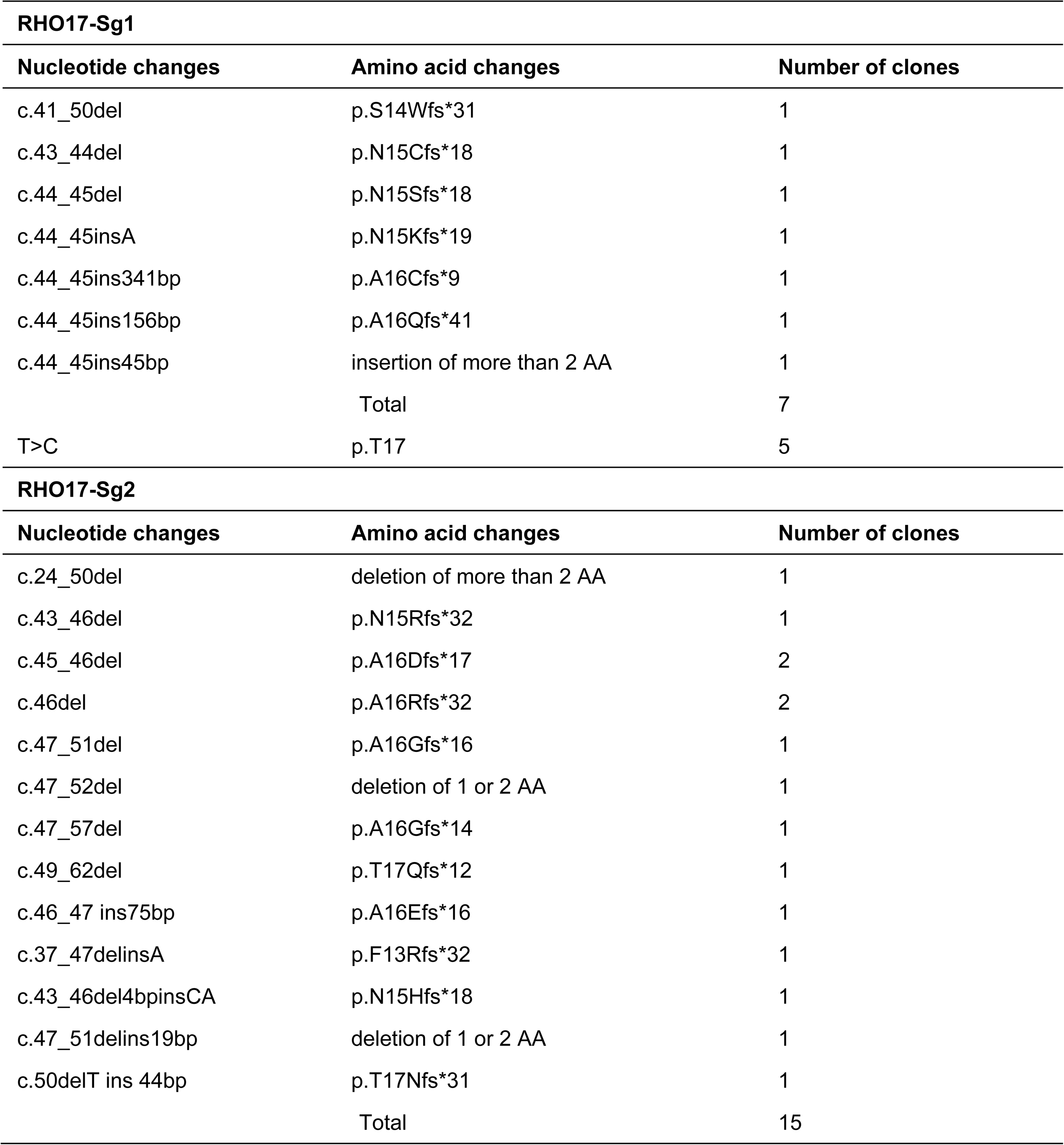
Results of TA cloning &Sanger Sequencing in 293T cells.

**Supplementary Table S2.**
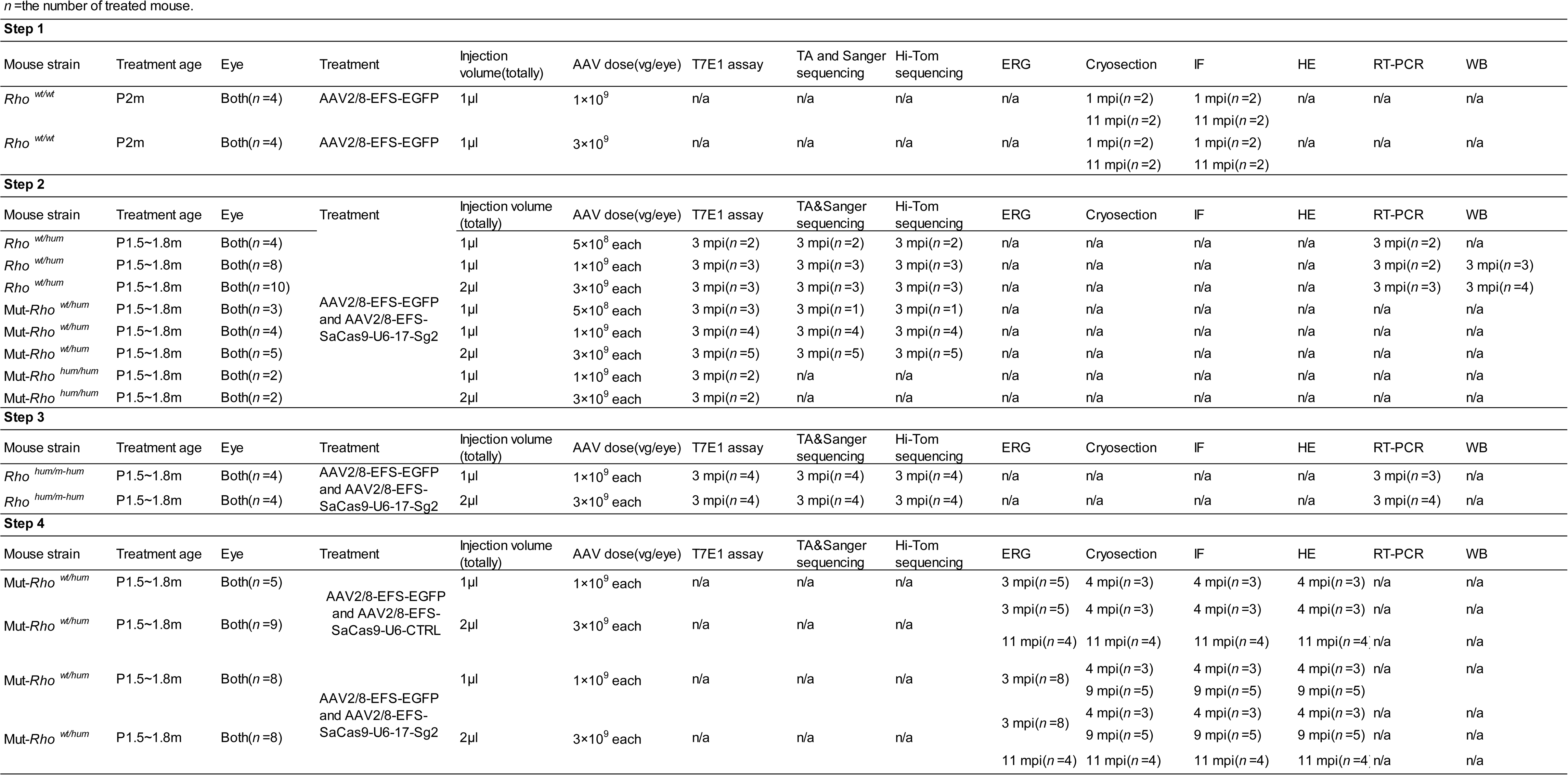
A comprehensive summary of mouse experiments.

**Supplementary Table S3.**
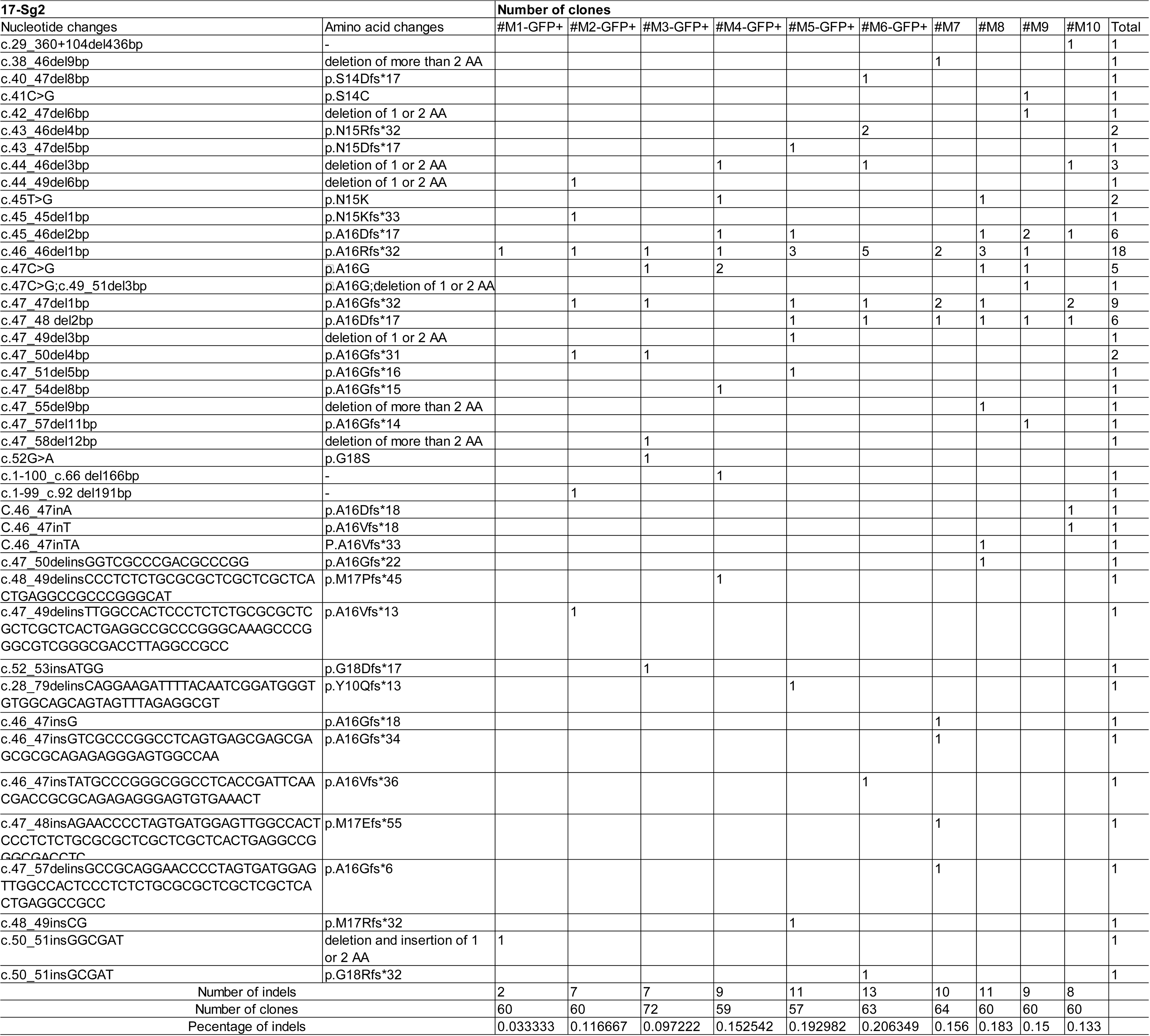
Results of TA cloning & Sanger sequencing in AAV-based SaCas9/17-Sg2-treated Mut-*Rho^wt/hum^* Retinas

**Supplementary Table S4.**
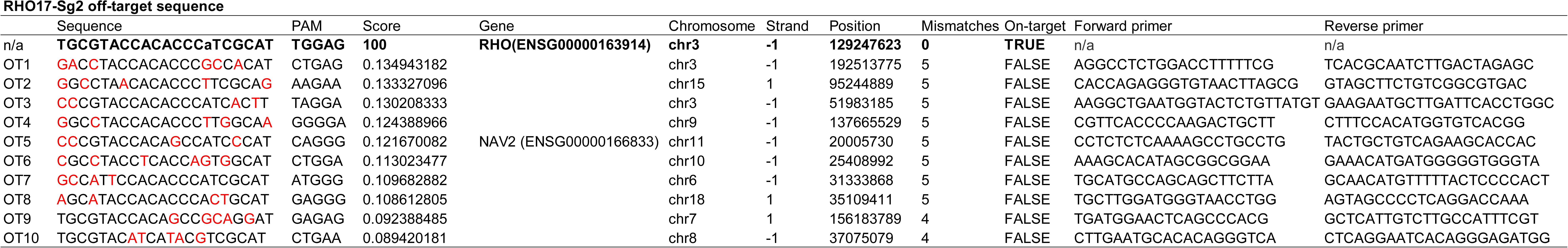
Off-target sites of 17-Sg2 obtained from Benchling (https://www.benchling.com/)

**Supplementary Table S5.**
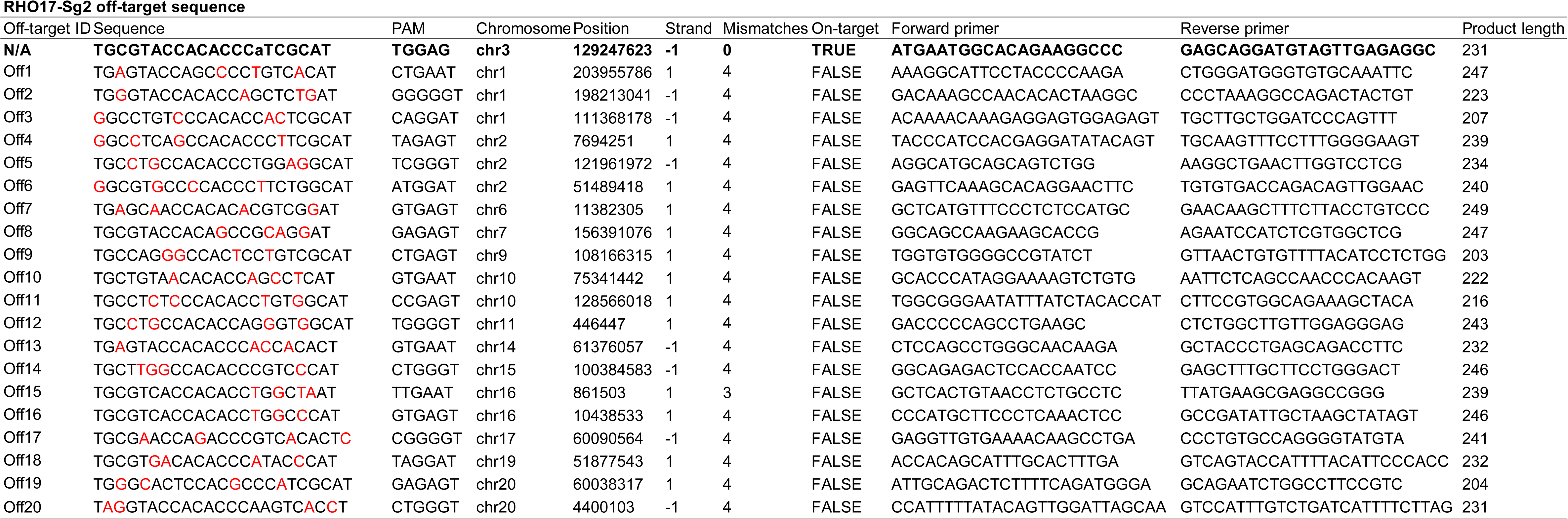
Off-target sites of 17-Sg2 obtained from Cas-OFFinder (http://www.rgenome.net/cas-offinder) and NGS results

**Supplementary Table S5.**
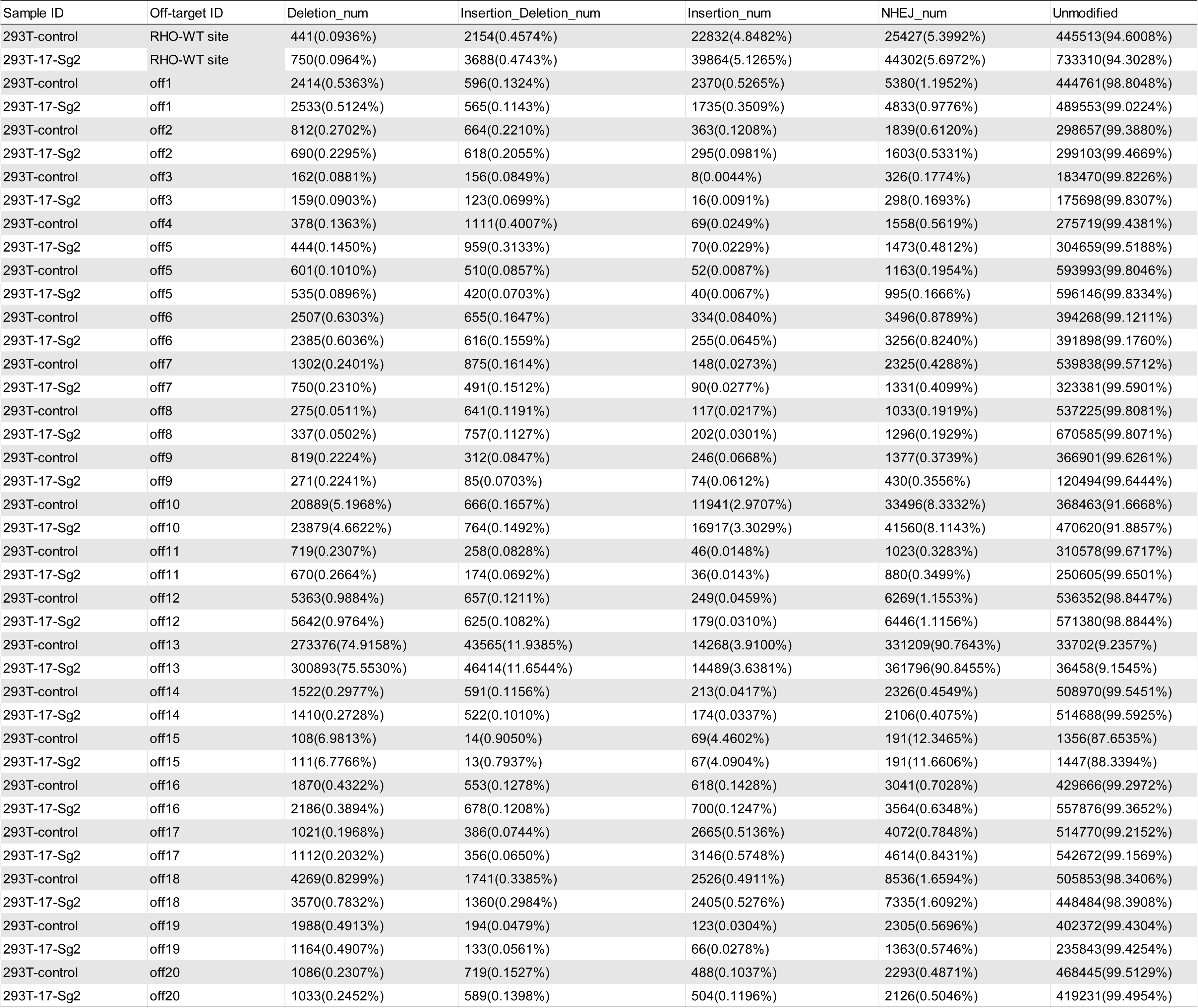
Off-target sites of 17-Sg2 obtained from Cas-OFFinder (http://www.rgenome.net/cas-offinder) and NGS results

**Supplementary Table S6.**
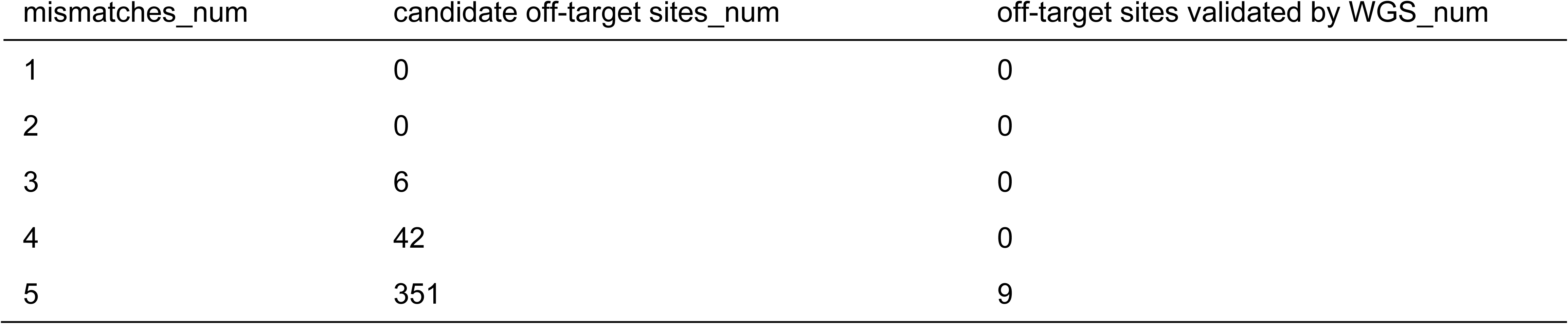
The number of off-target sites of 17-Sg2 obtained from Cas-OFFinder (http://www.rgenome.net/cas-offinder)

**Supplementary Table S7.**
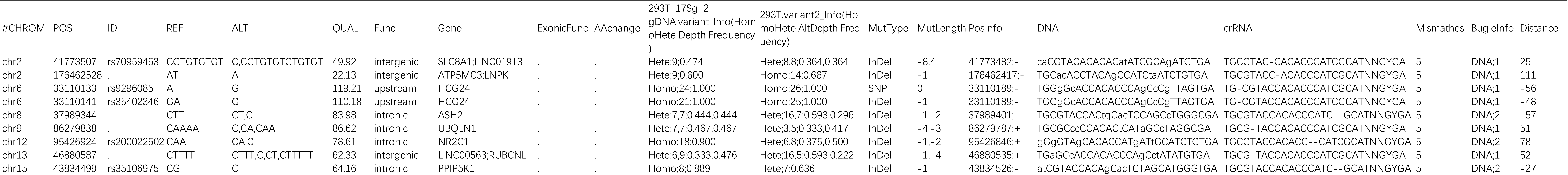
Whole-genome Sequencing results of off-target activity for SaCas9/17Sg-2.

**Supplementary Table S8.**
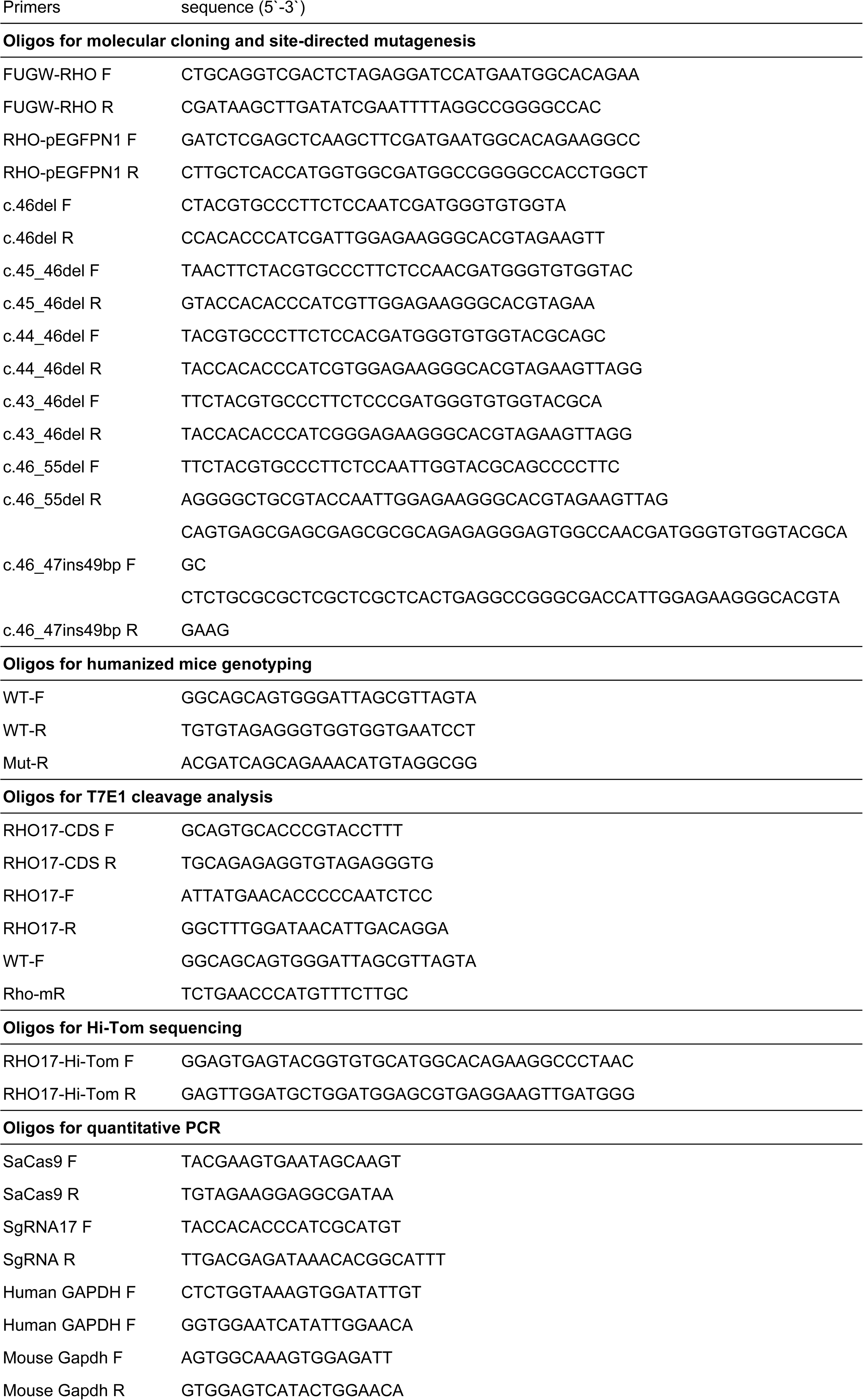
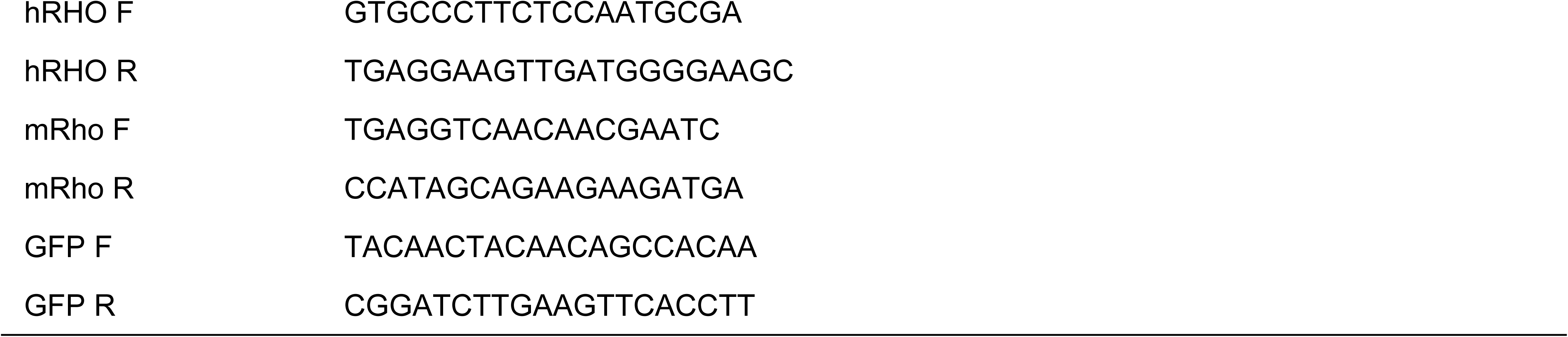
List of primers

